# Nuclear export inhibition jumbles epithelial-mesenchymal states and gives rise to migratory disorder in healthy epithelia

**DOI:** 10.1101/2022.06.11.495764

**Authors:** Carly M. Krull, Haiyi Li, Amit Pathak

**Author notes:** Corresponding author: Amit Pathak, Ph.D., One Brookings Dr., CB 1185, Saint Louis, MO 63130.

## Abstract

Epithelial-mesenchymal (E-M) phenotypes govern collective cellular behaviors to facilitate diverse tissue functions, including embryogenesis, wound healing, and cancer invasion. Cellular E-M state is regulated by dynamic nucleocytoplasmic transport of corresponding E-M factors; yet, it remains unknown how concurrently trapping these factors affects epithelia at the macroscale. To explore this question, we performed nuclear export inhibition (NEI) via Leptomycin B treatment, which biases nuclear localization of CRM1- associated E-M factors. We examined changes in collective cell migration across a range of substrate stiffnesses. Our results show that NEI fosters an atypical E-M state wherein cells concurrently strengthen intercellular adhesions and develop mechanoactive characteristics. Following NEI, soft substrates elevate collective migration for up to 24 h, while stiffer substrates reduce migration at all timepoints. We demonstrate that excluding Yes-associated protein 1 from NEI shifts affected cells toward an epithelial phenotype. Meanwhile, removing α-catenin maintains NEI’s intercellular adhesion strengthening and mechanoactivation capabilities, but prevents mechanoactive characteristics from reaching collective behavior. Overall, our results show that NEI disrupts epithelial migration through competition between intercellular adhesions, mechanoactivation, and cell-cell coordination. Ultimately, these findings of mechanoactive NEI outcomes for healthy cells could warrant additional investigation in the context of NEI-centered cancer therapies.

## Introduction

Collective cell migration facilitates diverse tissue functions, ranging from normal physiological activities, like embryogenesis and tissue repair, to malignant processes, like cancer invasion. While carrying out these functions, grouped cells can manifest diversely — as monolayers, clusters, or streams (Etienne-Manneville, 2014). Regardless of appearance, coordinated migration of the group is a carefully orchestrated balance between intercellular adhesive forces and intracellular propulsive ones (Trepat and Sahai, 2018). Cells transmit active forces from myosin molecular motors to their substrate (i.e., traction) to enable their forward motion. Meanwhile, those same motors generate tensile stress across adherens junctions, imparting strength to intercellular adhesions (Alert and Trepat, 2020). Cellular phenotypes arise, in part, from the degree of balance between these forces. When intercellular adhesion is high and anisotropic intracellular force is low, cells appear epithelial, presenting with apicobasal polarization and minimal motility (Yang et al., 2020). Reversal of that force balance gives rise to a mesenchymal phenotype; cells become more motile and invasive, polarizing front-back in their direction of migration. This shift in cellular characteristics constitutes a process termed epithelial-mesenchymal transition (EMT). However, far from a phenotypic switch, the transition between epithelial and mesenchymal states encompasses a spectrum of intermediate phenotypes, where a range of moderate intercellular adhesion and substrate tractions coincide. Ultimately, it is these intermediate states that enable the varying modes of collective cell migration.

Migrating epithelia comprise distinct leader and follower cell populations, which cooperate to facilitate group motion (Desai et al., 2013; Qin et al., 2021). Cells at the edge, the leaders, exhibit higher polarity and generate higher tractions (Reffay et al., 2011; Reffay et al., 2014; Trepat and Sahai, 2018). These cells apply most of the force required for locomotion, and they retain lower intercellular adhesion to do so (Matsuzawa et al., 2018). Meanwhile, cells behind – the followers – supplement leader tractions. Whereas followers maintain better adhesion to neighbors, they also extend cryptic protrusions to help forwardly propel the collective (Qin et al., 2021; Repat and Sahai, 2018). What arises from these distinct cell populations is an EMT gradient that fundamentally supports migration (Sarker et al., 2019). This gradient transpires from the tactful regulation of transcription factors (TFs) and migration-related proteins. Core EMT TFs, including Zeb, Snail, Slug, Twist directly regulate E-M phenotypes (Yang et al., 2020); and auxiliary migration-related proteins (e.g., YAP, IκBα, SOX9, and HIF2A, FOXA2) further contribute to movement between E-M states (Park et al., 2019; Huber et al., 2004; Huang et al., 2019; Yang et al., 2016; Zhang et al., 2015). However, while, on their own, these proteins unidirectionally affect cell fate (i.e., promote or inhibit EMT), it is unknown whether opposing factors compete, cooperate, or cancel during the construction of cell phenotypes.

To explore this question, we consider that the ability of EMT-TFs and related proteins to engender transcriptional changes hinges on their location within the cell: nuclear localization facilitates transcription of target genes, and cytoplasmic sequestration prevents it (Köster et al., 2005). Such trafficking between the nucleus and cytoplasm is termed nucleocytoplasmic (NC) transport (Cartwright and Helin, 2000). It denotes a highly controlled process, where nuclear import and export receptors ferry protein cargos through the nuclear pore complex to coordinate the signaling required for proper cell function. One way to survey interactions between opposing EMT factors is to force the nuclear co-localization of EMT-related proteins by biasing their import. Interfering with the appropriate nuclear export receptor achieves this outcome, and this process is called nuclear export inhibition (NEI). NEI has already been developed as a therapeutic strategy for cancer, where inhibition of the nuclear export receptor CRM1 returns critical tumor suppressor and oncogenic proteins to the nucleus and helps restore cell cycle regulation (Gravina et al., 2014; Gravina et al., 2017). However, CRM1-based NEI also biases the nuclear import of several key EMT-promoters (i.e., SNAIL, YAP, SOX9, HIF2A) and inhibitors (i.e., IκBα, FOXA2) (Xu et al., 2012). Here, we leverage this function of CRM1-based inhibitors to determine how nuclear co-localization of opposing EMT-related proteins manifests phenotypically. Interestingly, previous studies have found evidence only that CRM1-based NEI reverses EMT (Gravina et al., 2014; Azmi et al., 2015; Gravina et al., 2017; Kashyap et al., 2016a; Kashyap et al., 2016b; Galenski et al., 2021). However, E-M characteristics during NEI have been investigated thus far in a relatively limited context, discussed in terms of one or a few markers. Therefore, we wondered whether cellular outcomes could be more intricate than initially credited. To test this hypothesis, we treat monolayers composed of healthy MCF10A human mammary epithelial cells, chosen for their known E-M plasticity and mechanosensitivity, with the CRM1-based inhibitor of nuclear export, leptomycin B (LMB) (Kudo et al., 1999). We then assess changes in E-M cellular features and collective migration characteristics (Fig. 1).

**Figure 1.**
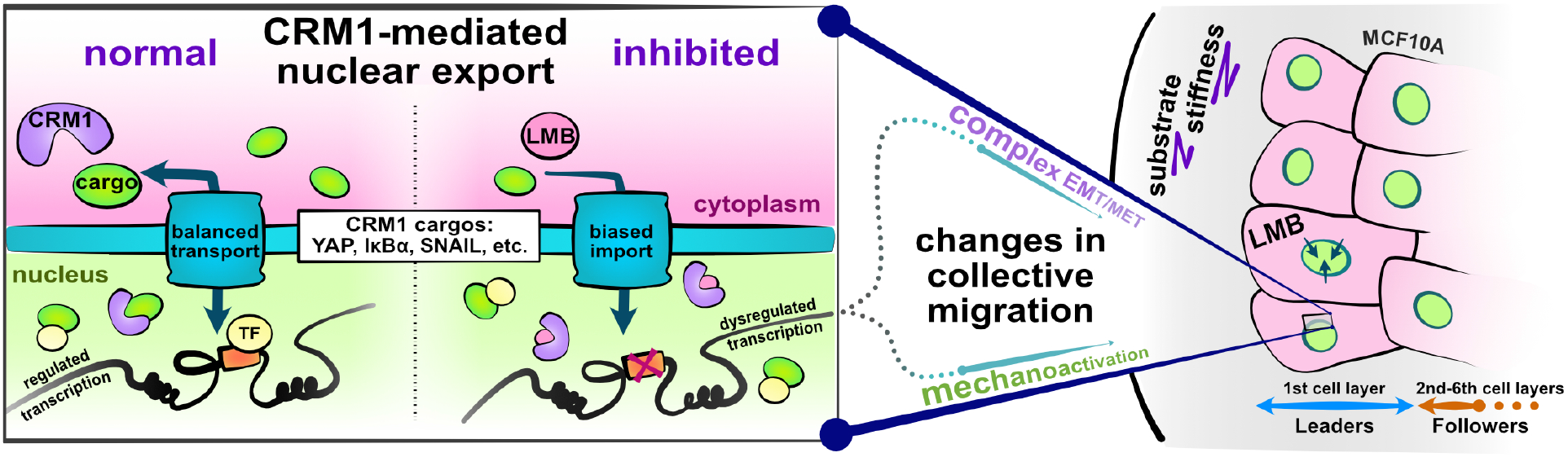
Schematic depicting experimental design and study hypotheses. The enlarged panel illustrates normal nucleocytoplasmic transport and accompanying well-regulated gene transcription (left). Changes following CRM1 inhibition (right) enable examination of the interaction between opposing EMT-related proteins in the development of collective cell phenotypes and migration characteristics. Depicted cells denote the experimental setup: MCF10A epithelial monolayers on polyacrylamide gels of varying stiffness, where leptomycin B is used to inhibit CRM1-mediated nuclear export.

Because this experimental framework depends greatly on NC transport, we also consider that the movement of proteins through nuclear pores is highly dynamic. Thus, the relative rates of import and export decide transcriptional outcomes. These rates are set by cargo size and geometry but are subject to change depending on the stiffness of the underlying substrate (Elosegui-Artola et al., 2017). More precisely, high stiffness promotes nuclear flattening and increases the nuclear import of proteins with higher molecular weight and stability. This process is one of the ways that stiffness-induced cellular mechanoactivation occurs, i.e., through Yes-associated protein 1 (YAP)-mediated signaling. Therefore, amidst competing EMT cues, substrate stiffness may contribute rate-dependent differences or additional mechanoactive features to phenotypic outcomes (Walter et al., 2018; Fattet et al., 2020; Wei et al., 2015). To uncover this potential stiffness-dependency, we compare NEI effects for MCF10A epithelia seeded on polyacrylamide gels spanning a range of stiffnesses.

Given the complexity of EMT signaling arising from the combined NEI and substrates cues, in this study we describe epithelial and mesenchymal cellular traits according to their physical characteristics, migratory phenotypes, and candidate protein expressions, but not in terms of selected gene activations, as discussed previously (Yang et al., 2020). Ultimately, we find that NEI forces competition between cellular E-M states, which generally leads to disordered collective migration; yet, whether NEI promotes or inhibits migration depends distinctly on the underlying matrix stiffness. Together, these experiments demonstrate that nuclear export inhibition traverses length scales – concurrently trapping E-M factors, jumbling cellular E-M states, and disrupting grouped epithelial migration. Overall, the presented work may have implications for cancer, wound healing, and embryonic development.

## Results

### NEI causes preferential nuclear localization of IκBα and strengthens cell-cell adhesions in a stiffness-dependent manner

Previous investigations have proposed that NEI reverses EMT, and mechanistic studies indicate this reversal may transpire via IκBα-dependent inhibition of NFκB (Fig. 2A) (Gravina et al., 2014; Azmi et al., 2015; Gravina et al., 2017; Kashyap et al., 2016a; Kashyap et al., 2016b; Galenski et al., 2021). However, E-M characteristics during NEI have been investigated thus far in a relatively limited context, discussed in terms of one or a few markers, and with culture performed primarily on tissue culture plastic. Because NEI causes nuclear localization of multiple competing EMT-related proteins and stiff substrates themselves trigger EMT through mechanoactive signaling, we wondered whether the manner in which NEI changes cellular E-M state could be more intricate than initially credited. To investigate this question, we perform a broad analysis of cell phenotype across mechanically varying substrates. Here, we begin by assessing IκBα localization and intercellular adhesion strength to determine how NEI influences epithelial characteristics in our model system.

**Figure 2.**
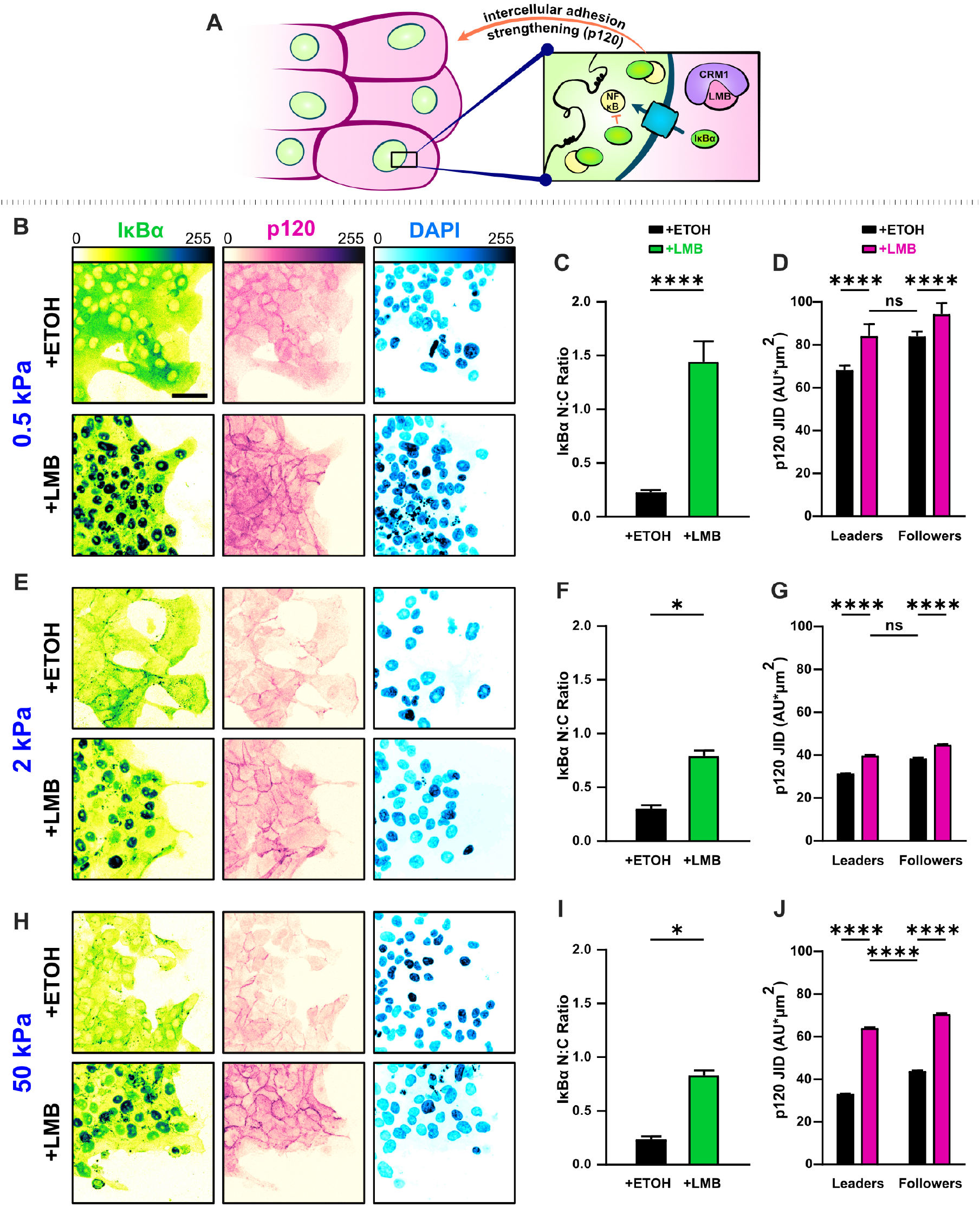
NEI enriches epithelial features in MCF10A collectives. **(A)** Schematic illustrating the established relationship between LMB and cell-cell adhesion strengthening, where nuclear accumulation of IκBα promotes inhibition of NF-κB. **(B, E, H)** Representative images for epithelial characteristics of WT MCF10A on 0.5, 2, and 50 kPa polyacrylamide gels, respectively. Monolayers were treated with ethanol (ETOH) as vehicle or leptomycin B (LMB) for NEI. Images depict nucleocytoplasmic localization of IκBα (left), p120 expression (middle), and DAPI nuclear signal (right). **(C, F, I)** Nucleocytoplasmic (N:C) ratio for IκBα (n = 8). Data was analyzed using a two-way ANOVA to evaluate NEI and stiffness effects. **(D, G, J)** Leader-follower changes in p120 junction integrated density (JID) (n > 39 leaders and 55 followers). Data was analyzed using a three-way ANOVA with Tukey post hoc analyses to evaluate NEI, stiffness, and leader-follower differences. Bars represent mean ± SEM. Scale bar: 50 µm.

Nuclear localization of IκBα is an established measure for NFκB inhibition (Arenzana-Seisdedos et al., 1997), and junctional expression of p120 reflects the degree of cell-cell adhesion reinforcement (Kourtidis et al., 2013). Therefore, to determine whether epithelial characteristics changed with substrate stiffness, we surveyed the localization of IκBα and junctional p120 levels on 0.5, 2, and 50 kPa polyacrylamide gel. Consistent with prior studies in cancer cells, we found that NEI contributed to higher IκBα nucleocytoplasmic (N:C) ratios (Fig. 2B-C, E-F, H-I). The preferential nuclear localization of IκBα appeared in both leader and follower cells. However, the strength of epithelial characteristics did depend on substrate stiffness. Nuclear localization of IκBα was higher on 0.5 kPa than on 2 and 50 kPa, and junctional p120 levels exhibited a matching stiffness effect (Fig. 2D, G, J). Yet remarkably, across stiffnesses, the conventional relationship between leader and follower cells – where followers adhere more strongly to their neighbors – was preserved. Together, these findings suggest that mechanoactivation, as translated through substrate stiffness, limits NEI’s ability to inhibit NFκB and reinforce cell-cell adhesions. Despite this, NEI upholds traditional leader-follower cell-cell adhesion gradients independent of substrate stiffness.

### Mechanoactivation during NEI evolves together with reinforced intercellular adhesion to promote a concurrent epithelial-mesenchymal state

Because NEI causes nuclear accumulation of the mechanoactivating protein YAP, together with multiple EMT-promoting proteins (i.e., SNAIL, SOX9, HIF2A), we wondered whether mechanoactive cellular characteristics would develop (Fig. 3A). To answer this question, we first validated that nuclear localization of YAP occurs during NEI across substrate stiffnesses (Fig. 3B,C). Confirmation of elevated nuclear YAP signified that mechanoactivation might be present independent of substrate triggers. Interestingly, we also found that the shift toward nuclear localization of YAP occurred in both leader and follower cells. Thus, while NEI preserved the intercellular adhesion gradients customary to leader and follower populations, the associated mechanoactivation gradients dissolved.

**Figure 3.**
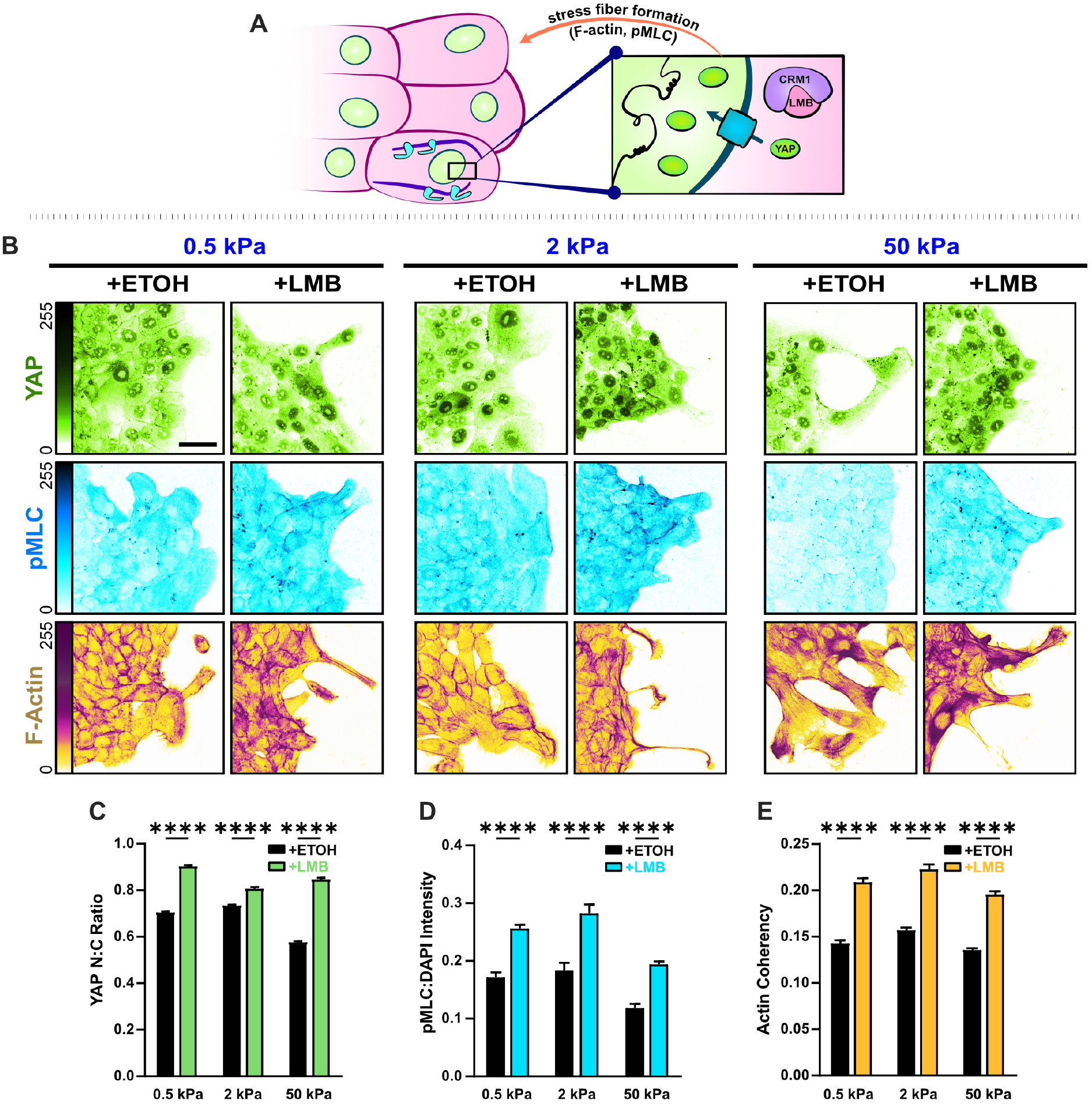
Mechanoactive and mesenchymal features develop in concert with epithelial characteristics during NEI. **(A)** Schematic illustrating the established relationship between YAP activation and stress fiber formation. **(B)** Representative images for mesenchymal characteristics of WT MCF10A on 0.5, 2, and 50 kPa polyacrylamide gels. Monolayers were treated with ethanol (ETOH) as vehicle or leptomycin B (LMB) for NEI. Images depict nucleocytoplasmic localization of YAP (top), pMLC expression (middle), and F-actin (bottom). **(C)** Nucleocytoplasmic (N:C) ratio for YAP (n > 55 leaders and 220 followers), **(D)** pMLC intensity (n = 8), and **(E)** actin coherency (n > 1000) for all stiffnesses. Data was analyzed using a two-way ANOVA with Tukey post hoc analyses to evaluate NEI and stiffness effects. Bars represent mean ± SEM. Scale bar: 50 µm.

We considered how this might change cell morphology, pMLC expression, and F-actin coherency. During migration, we found, with high frequency, that NEI caused leader cells to protrude as thin extensions from the monolayer baseline (Fig. 3B). Comparatively, control leaders maintained more conventional fan-like shapes, even at high stiffnesses. Contributing to these morphological differences, the actin coherency of protruding cells was higher following NEI (Fig. 3E); meanwhile, pMLC expression was elevated universally in leaders and followers (Fig. 3D). Curiously, these results suggest that epithelial characteristics (e.g., reinforced intercellular adhesions) and mesenchymal features (e.g., higher cytoskeletal forces, front-back polarization) evolve together during NEI.

### NEI invariably leads to migratory disorder, but changes in migration speed depend on substrate stiffness and adhesion reinforcement

Because epithelial cells balance intercellular adhesion and mechanoactivation gradients to facilitate collective movement, we hypothesized that the observed disturbances could interfere with the typical patterns of migration. To assess these potential migration changes, we collected time-lapse images of migrating epithelia for 24 h, then used particle image velocimetry to determine how NEI affects conventional patterns of migration.

Results reveal stiffness-dependent changes in migration speed, velocity, and order. Previous studies have exclusively shown that NEI inhibits migration; yet, strikingly, on 0.5 kPa, NEI elevated collective migration velocity (Fig. 4A, D). Higher cellular speeds within the monolayer facilitated this elevation of net migration (Fig. 4G). Meanwhile, the rise of cell backtracking and overall disorder contributed to the decrease of net velocity toward vehicle levels, beginning around 3 h (Fig. 4H, M).

**Figure 4.**
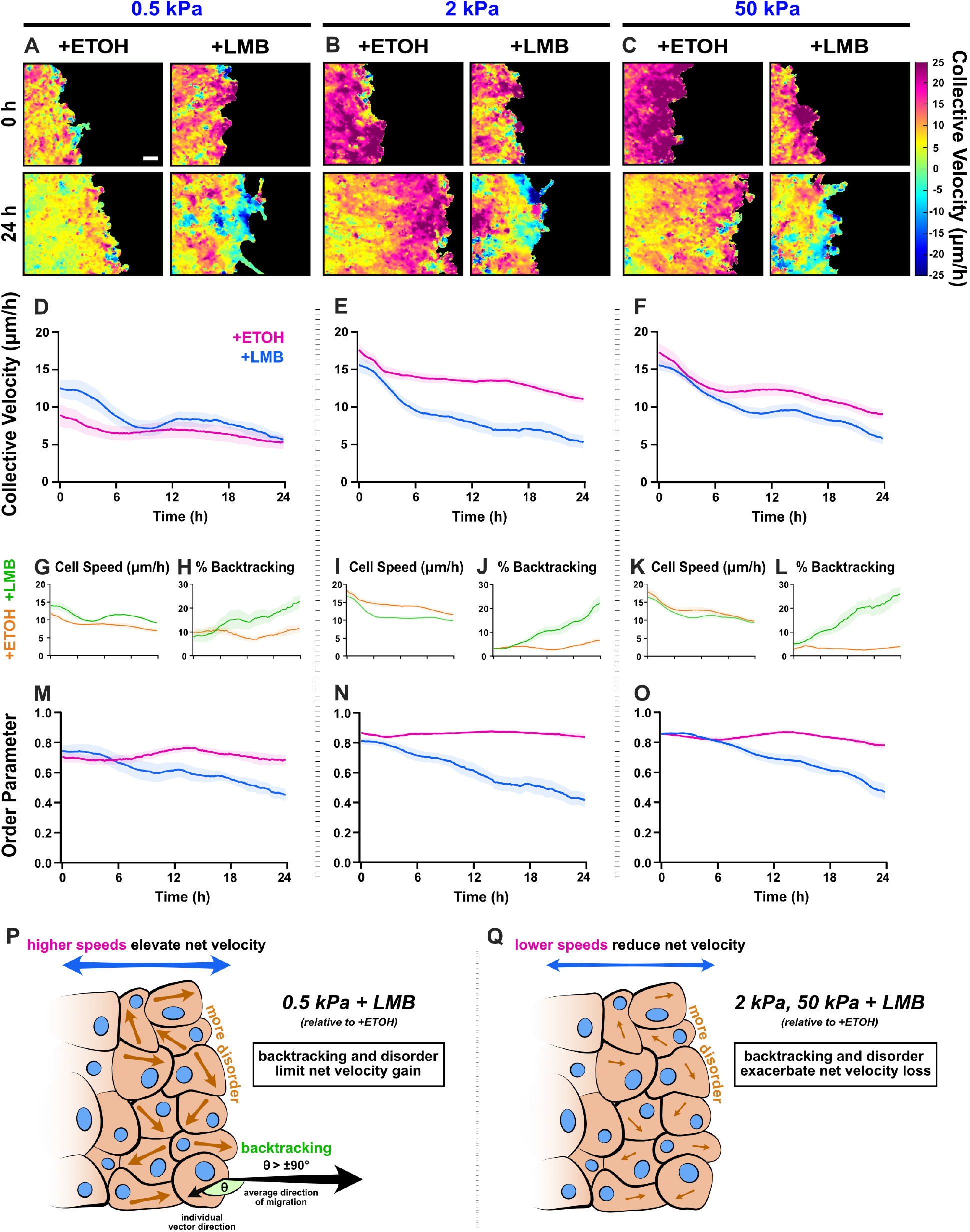
Collective migration changes from NEI are stiffness-dependent. Velocity heat maps generated from particle image velocimetry analysis of 24 h migration for **(A)** 0.5 kPa, **(B)** 2 kPa, and **(C)** 50 kPa, (n ≥ 6). **(D-F)** Respective quantifications of net velocity over time. Average speeds for **(G)** 0.5 kPa, **(I)** 2 kPa, and **(K)** 50 kPa, along with the % of backtracking vectors **(H, J, L)**, respectively. Order parameter for **(M)** 0.5 kPa, **(N)** 2 kPa, and **(O)** 50 kPa. Schematics describing how NEI changes migration characteristics for (**P)** 0.5 kPa and **(Q)** 2 and 50 kPa. Lines represent mean ± SEM. Scale bar: 100 µm.

In contrast, on 2 and 50 kPa, NEI decreased net migration velocity at all time points (Fig. 4E, F). For 2 kPa, this decrease was mediated in initial stages by reduced cellular speeds (Fig. 4I). Meanwhile, the progressive loss of order and associated increase in cell backtracking exacerbated the overall velocity loss with time (Fig. 4J, N). For 50 kPa, early time points (0-6 h) showed only slight decreases in net migration velocity (Fig. 4F). Higher cell backtracking (Fig. 4L) accounted primarily for the net migration slowing, since cell speeds were only marginally lower compared to vehicle control (Fig. 4K), and order was temporarily preserved (Fig. 4O). Across time points, cell speeds on 50 kPa were comparable to vehicle. Therefore, 50 kPa results resemble 0.5 kPa, where, although speeds were higher, they remained similar to vehicle control. Overall, despite only marginal losses in cell speed on 50 kPa, backtracking together with rising disorder facilitates the observed decrease in collective velocity. Taken as a whole, these findings depict changes in the conventional patterns of collective migration during NEI, where disorder and cell backtracking escalate, independent of substrate stiffness. These results further uncover stiffness-dependent NEI outcomes, where, counter to reports of migration inhibition exclusively, NEI promotes migration on soft substrates.

### Disruption of Golgi polarization in leaders and global elevation of gm130 expression underlie NEI-induced disorder

The Golgi apparatus contributes to front-rear polarization by orienting in front of the nucleus in the direction of migration (Ravichandran et al., 2020). Moreover, the associated transport of proteins and nucleation of microtubules toward the front of migrating cells supports directed motion (Yadav et al., 2009). To determine whether changes in the Golgi influenced rising disorder during NEI, we analyzed gm130, a Golgi marker that assists microtubule nucleation and the development of mesenchymal cellular phenotypes (Rivero et al., 2009; Baschieri et al., 2014). First, we assessed whether orientation of the Golgi changed in leader and follower cells. To do this, we defined a polarized Golgi as one where the Golgi centroid resided within 60° of the direction of migration, when measured from the associated nuclear centroid (Fig. 5A) (Mason et al., 2019). For epithelia treated with vehicle, we found that stiffer substrates promoted leader cell Golgi polarization. Almost twice as many leaders displayed a polarized Golgi apparatus at 50 kPa compared to 0.5 kPa (Fig. 5B,C). This is consistent with stiffness-induced cell polarization, which has been reported previously (Alert and Trepat, 2020). Meanwhile, followers treated with vehicle exhibited a biphasic polarization curve, where 2 kPa had the highest percentage of polarized cells (Fig. 5D).

**Figure 5.**
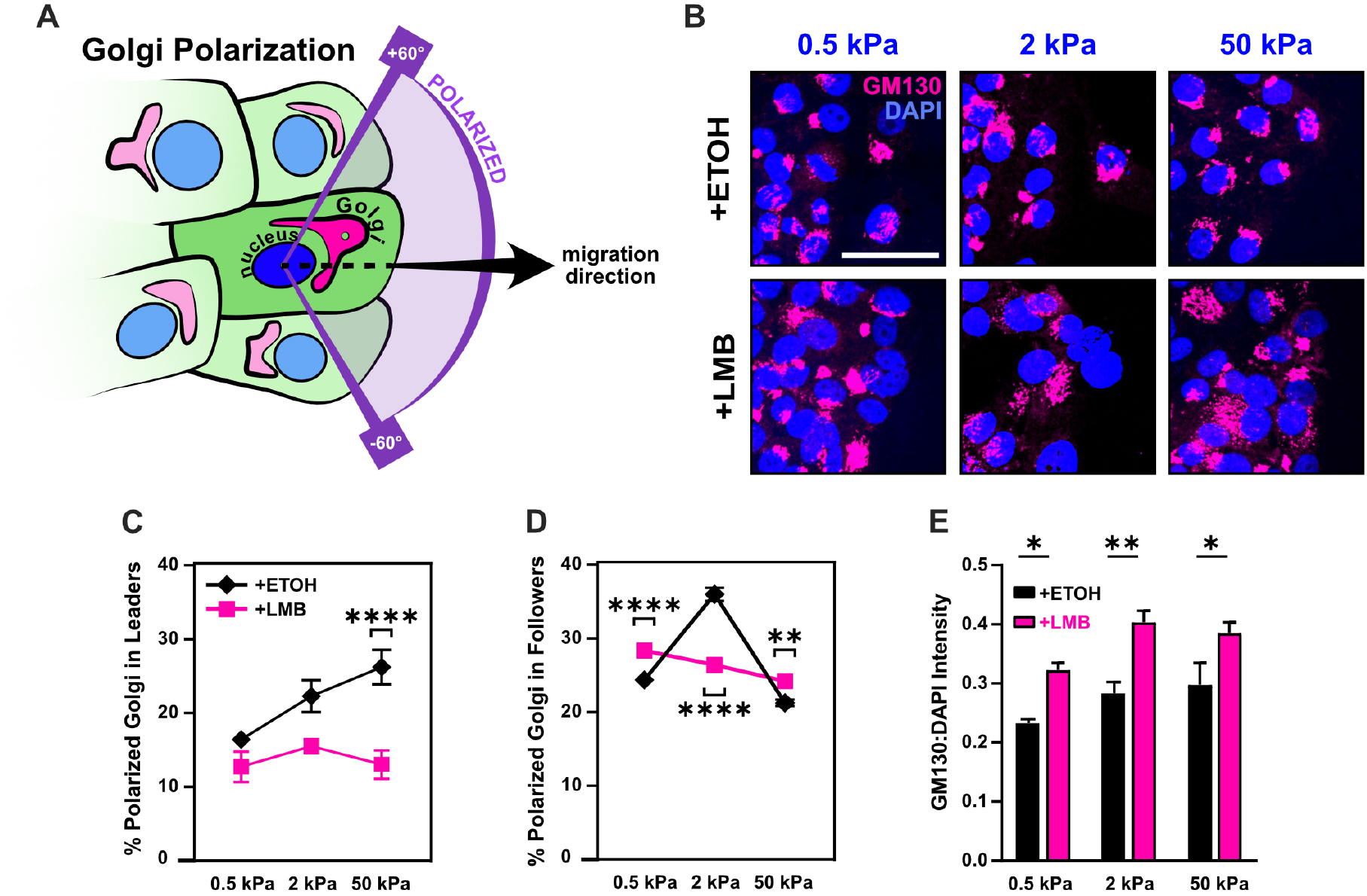
Disrupted Golgi polarization and GM130 expression underlie disordered migration following NEI. **(A)** Schematic definition for Golgi polarization, where a polarized Golgi is one whose centroid lies within 60° of the line drawn from the nuclear centroid in the average direction of migration. **(B)** Representative images for gm130 and DAPI across substrate stiffnesses and cell treatments. Quantification of Golgi polarization in **(C)** leaders (n > 50) and **(D)** followers (n > 250). Shapes represent mean and error bars represent SEM. **(E)** Quantification of gm130 intensity across substrate stiffnesses (n = 8). Bars represent mean ± SEM. All data was analyzed using two-way ANOVAs with Tukey post hoc analyses to evaluate NEI and stiffness effects. Scale bar: 50 µm.

NEI’s most prominent effect on Golgi orientation was in leader cells. While leader cells customarily provide directional cues for followers, following NEI, stiffness-dependent polarization of leaders disappeared. Moreover, leader polarization percentages dropped below those reported for control cells. This finding was consistent with the cyclic protrusion and retraction of leader cells observed during migration with NEI and presumably contributed to leading-edge instability. It also reflects previous reports where the rearward position of the Golgi emerged after transition to a mesenchymal cellular state (Natividad et al., 2018). Meanwhile, the effect of NEI on follower polarization was more subtle and changed with substrate stiffness. On 0.5 and 50 kPa, NEI increased follower polarization. This increased Golgi polarization appropriately corresponded to the higher junctional stability, velocities, speeds, order, and backtracking observed during migration. Congruently, the lower polarization on 2 kPa corresponded to reduced migration parameters (Fig. 4). However, while NEI’s impact on Golgi orientation changed with substrate stiffness, its effect on gm130 expression itself was consistent – displaying increased levels across stiffnesses (Fig. 5E). Together, these results suggest that, in this system, gm130 expression may support the cytoskeletal rearrangement required for the front-back mesenchymal-like polarization observed in leaders, and Golgi orientation itself may contribute to the loss of order observed during NEI.

### Leading-edge instability and combined epithelial-mesenchymal characteristics promote multicellular stream formation during NEI

Previous studies established that leading-edge instability, higher intercellular adhesion, localized mechanoactivation, and coordinated cellular velocities together support the formation of multicellular streams (Vishwakarma et al., 2018; Sarker et al., 2019). Here, we have shown that NEI destabilizes the leading edge, demonstrated via decreased leader polarization; reinforced intercellular adhesion, measured via increased junctional p120; caused mechanoactivation, evidenced via higher YAP N:C ratio, pMLC expression, and F-actin coherency; and contributed invasive characteristics, shown through elevated gm130 expression. Therefore, despite the reduced overall coordination of velocity during NEI, reflected in low order parameter, we hypothesized that, at some level, NEI might encourage multicellular streaming (Fig. 6A). To investigate this hypothesis, as previously, we defined a cell stream as a protrusion of 2 or more cells in succession from the monolayer baseline, such that protrusion length was more than twice its width (Fig. 6B) (Sarker et al., 2019). After examining migration time-lapses, we found that transient cell streams formed, even during migration with vehicle (Fig. 6C). However, these streams formed at relatively low frequency (Fig. 6D) and with few cells (Fig. 6E). The presence of streams in this control condition suggests that fluctuations in intercellular adhesion and mechanoactivation at the leading edge were sufficient to form small, transient streams. By contrast, streams during NEI were more exaggerated (Fig. 6C, Fig. 4A-C), evolving at higher frequency and with more cells in each stream (Fig. 6D-E). Higher stream formation during NEI thus indicates that adhesion strength and mechanoactivation may stimulate brief periods of cell streaming, even in regions of high disorder. Therefore, while previous studies show that structural changes in the underlying matrix (e.g., collagen fiber orientation, topography) trigger cell streaming via adhesion-based signaling and consequent mechanoactive protein expression, these results highlight that streaming can also arise from direct modifications to nuclear export.

**Figure 6.**
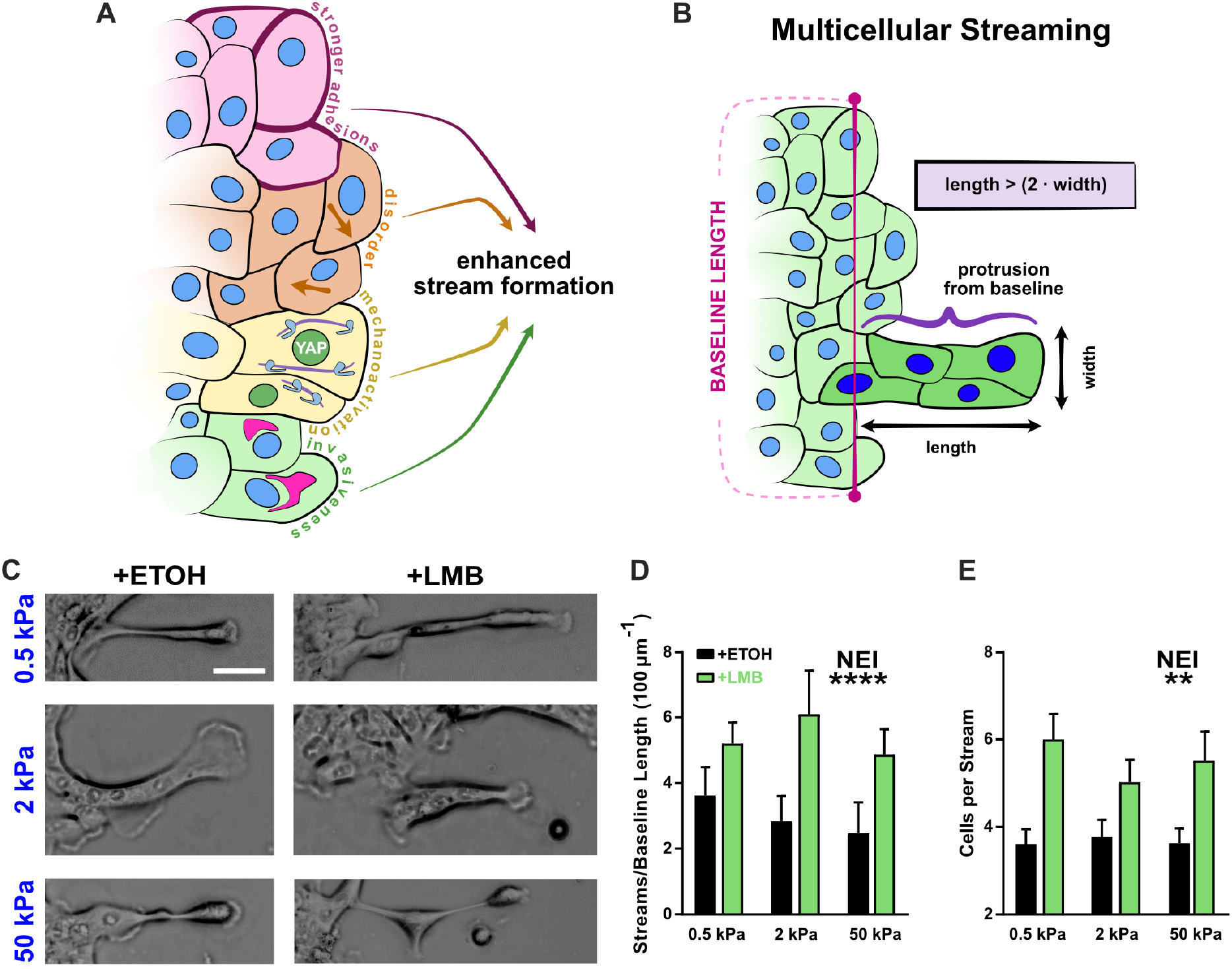
Concurrent epithelial-mesenchymal features support multicellular streaming. **(A)** Schematic depicting NEI-induced changes in cell phenotype and migration, which imply the potential for enhanced stream formation. **(B)** Schematic definition for multicellular streams. **(C)** Example images for multicellular streams seen across stiffnesses and treatment conditions. **(D)** Accompanying quantification for the number of **(D)** streams per baseline length and **(E)** cells per stream. Data was analyzed using two-way ANOVAs with Tukey post hoc analyses to evaluate NEI and stiffness effects. *’s denote the significance level for NEI effects. Scale bar: 50 µm.

### YAP silencing shifts NEI-affected cells toward an epithelial state by strengthening intercellular adhesion and attenuating mechanoactivation

NEI causes nuclear localization of YAP, which leads to the acquisition of mechanoactive phenotypes, such as that previously shown (Fig. 3). Given its role as an EMT initiator (Park et al., 2019), we hypothesized that excluding YAP from the affected cargos could reduce mesenchymal characteristics and shift cells toward a more conventional epithelial state. We examined the effect of YAP silencing across 0.5, 2, and 50 kPa stiffnesses (Fig. S1-S3). However, to discuss the core differences observed during NEI without YAP, we focus on results from the intermediate stiffness, 2 kPa (Fig. 7).

**Figure 7.**
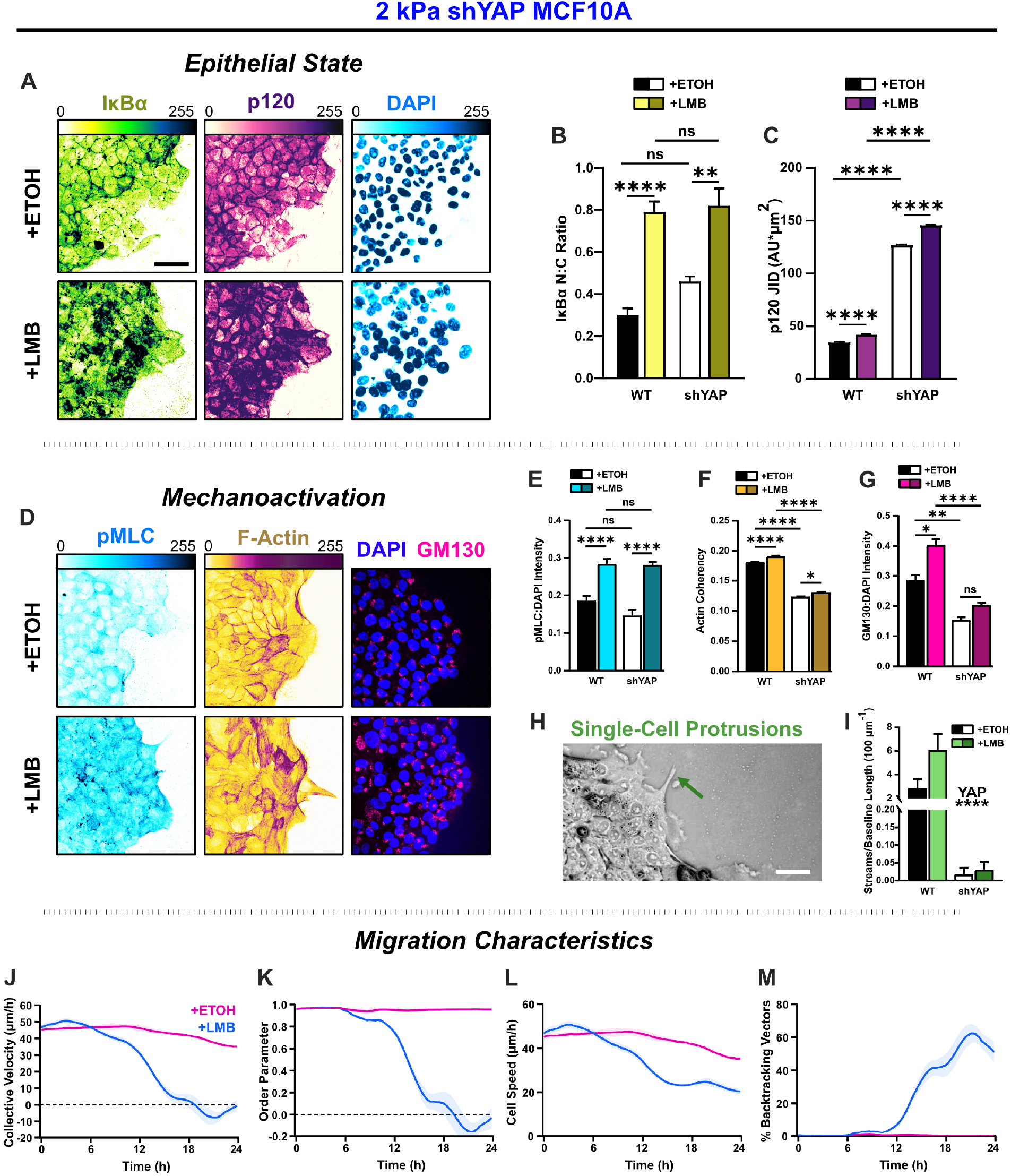
YAP silencing enhances epithelial characteristics and attenuates mechanoactivation. **(A)** Epithelial characteristics for shYAP MCF10A. Images depict nucleocytoplasmic localization of IκBα (left), p120 expression (middle), and DAPI nuclear signal (right). **(B)** Nucleocytoplasmic (N:C) ratio for IκBα (n = 8). **(C)** Changes in p120 junction integrated density (JID) (n = 8). **(D)** Mechanoactive characteristics for shYAP MCF10A. Images depict pMLC (left), F-actin (middle), and gm130 expression (right). Quantification comparing WT and shYAP **(E)** pMLC expression (n = 8), **(F)** actin coherency (n > 1000), and **(G)** gm130 expression (n = 8). **(H)** Representative image showing single-cell protrusions at the leading edge. **(I)** Quantification of multicellular streams per baseline length for WT and shYAP cells (n ≥ 6). Bars represent mean ± SEM. All data was analyzed using two-way ANOVAs with Tukey post hoc analyses to evaluate NEI and YAP effects. For streams, *’s denote the significance level for YAP effects. Time plots of migration characteristics (n ≥ 6): **(J)** net velocity, **(K)** order parameter, **(L)** speed, and **(M)** percentage of backtracking vectors for shYAP monolayers on 2 kPa polyacrylamide gel. Lines represent mean ± SEM. Scale bars: 50 µm.

In contrast to WT cells, where IκBα localized cleanly within the cytoplasm or nucleus, shYAP cells exhibited punctate staining at intercellular adhesions in control conditions and displayed strong cytoplasmic signals during NEI. These differences are consistent with IκBα degradation, a known by-product of YAP silencing (Wang et al., 2020). However, YAP silencing significantly increased p120 expression, indicating stronger intercellular adhesion than WT for all conditions (Fig. 7A-C). This finding suggests that removing YAP from the affected cargos enhances epithelial aspects of NEI’s concurrent E-M phenotype.

To determine how YAP silencing affected mesenchymal characteristics, we next examined differences in pMLC, F-actin, gm130, and cell morphology during NEI. Interestingly, pMLC levels after NEI were unchanged with YAP silencing (Fig. 7D-E). However, YAP-silenced cells exhibited reduced actin coherency and gm130 expression relative to WT cells (Fig. 7F-G). Meanwhile, NEI-related changes in gm130 expression were insignificant. Consistent with this reduction in the cytoskeletal features required for mesenchymal presentation, YAP-silenced leaders formed much shorter protrusions at the leading edge (Fig. 7H), and multicellular streams were also inhibited (Fig. 7I). Together, these results demonstrate that isolating YAP from affected cargos attenuates mesenchymal aspects of the combined E-M phenotype induced during NEI.

This shift in E-M state led to significant changes in collective migration. For NEI on 2 kPa, YAP-silenced cells exhibited sudden drops in net migration velocity beginning around 12 h. Migration on 0.5 and 50 kPa exhibited similar drops, which preceded the complete arrest of net migration for all stiffnesses (Fig. S3 A-C). Moreover, at later time points, cells transiently reversed their direction of migration (Fig. 7J, Fig. S3 A-C). Results suggest that changes in migration order prompted those changes in velocity. At 12 h, the sudden drop in net velocity coincided with a sudden drop in migration order, and at later time points, migration velocity and order parameter jointly oscillated about zero (Fig. 7J-K). Interestingly, and contrary to WT, YAP-silenced cells exhibited non-monotonic increases in cell backtracking (Fig. S3 E, G, I). Inflection points for backtracking coincided with inflection points for net velocity, order parameter, and speed. These points were present across substrate stiffnesses and reflected transient oscillations in order within the monolayer (Fig. S3). In general, the observed worsening of net velocity and order losses during NEI is consistent with previous findings of impersistent cell motility following YAP depletion (Mason et al., 2019). Therefore, while YAP contributes to mechanoactive features during NEI, it also provides crucial directional cues to assist collective migration. In other words, YAP may enhance cohesion to temper the loss of order during NEI.

Notably, NEI initially increased shYAP cell speeds on 0.5 and 2 kPa, which could indicate residual YAP-independent mechanoactivation. However, after 6 h, speeds were lower than vehicle control for all substrate stiffnesses (Fig. S3 D, F, H). This time-dependency could indicate that early NEI changes are dominated by mechanoactive behavior, while cell-cell adhesion changes occur later. Overall, these results demonstrate that YAP silencing shifted NEI’s concurrent E-M state by enhancing epithelial aspects and attenuating mesenchymal ones.

### α-catenin knockdown shifts NEI-affected cells toward a mesenchymal state but prevents the transfer of mechanoactive features to collective behaviors

Propagation of forces through adherens junctions is an element fundamental to collective migration (Ladoux and Mège, 2017). One of the central mediators in this cooperative force propagation serving to mechanically couple cells is the protein α-catenin, which recruits F-actin to E-cadherin-based cell junctions. Therefore, to characterize the role of intercellular force propagation in the observed NEI outcomes, we used α-catenin depleted MCF10A cells, as described previously (Loza et al., 2016). We examined the effect of α-catenin knockdown across 0.5, 2, and 50 kPa stiffnesses (Fig. S4-S6). However, to discuss the key outcomes from α-catenin depletion, we again focus on results from the intermediate stiffness, 2 kPa (Fig. 8).

**Figure 8.**
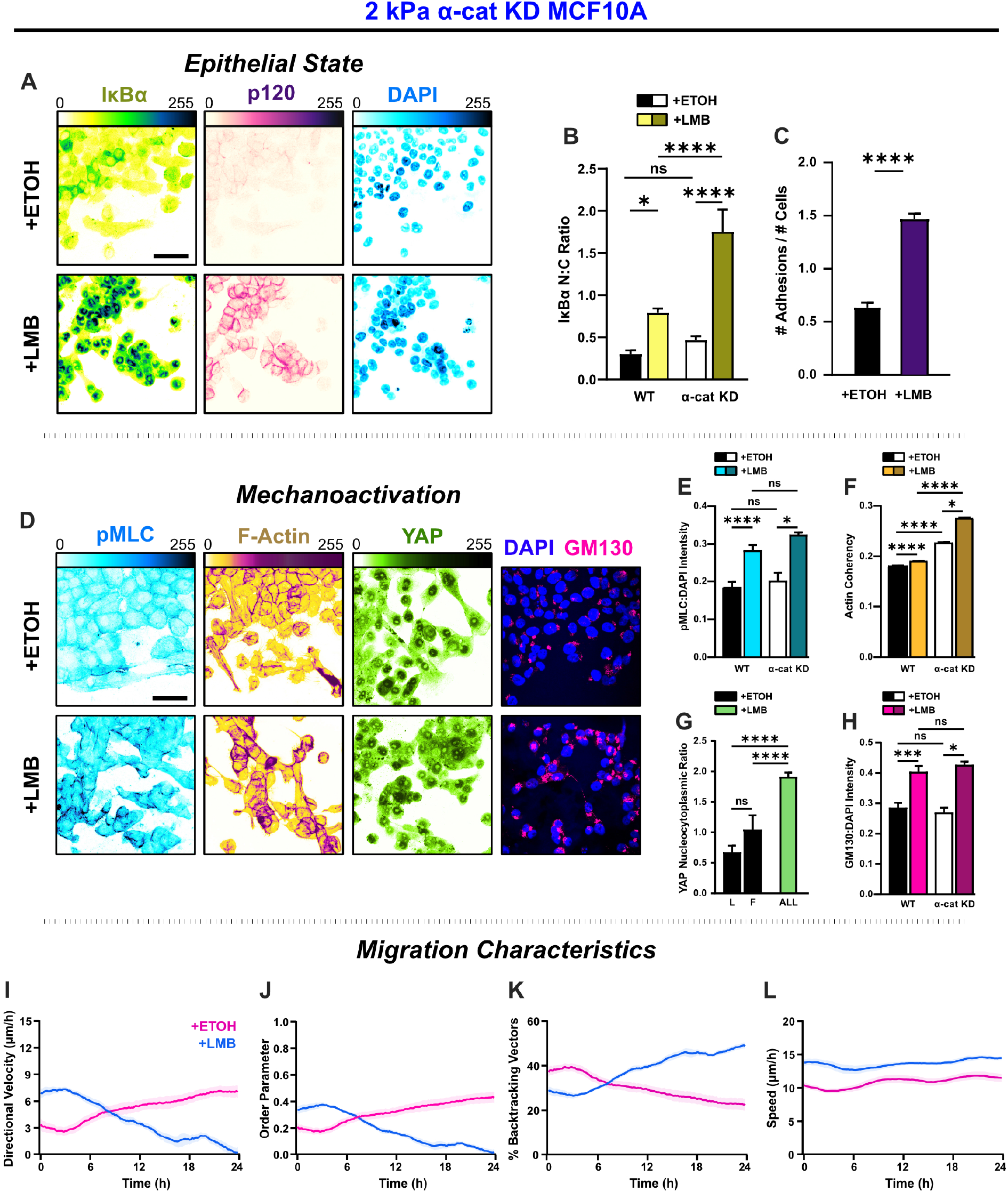
α-catenin knockdown prevents the transfer of mechanoactive features to collective cell behavior. **(A)** Epithelial characteristics for α-cat KD MCF10A. Images depict nucleocytoplasmic localization of IκBα (left), p120 expression (middle), and DAPI nuclear signal (right). **(B)** Nucleocytoplasmic (N:C) ratio for IκBα (n = 8). **(C)** Changes in the number of discernible p120-marked junctions (n = 8). **(D)** Mechanoactive characteristics for α-cat KD MCF10A. Images depict pMLC (left), F-actin (middle), and gm130 expression (right). Quantification comparing WT and α-cat KD **(E)** pMLC expression (n = 8), **(F)** actin coherency (n > 1000), **(G)** nucleocytoplasmic (N:C) YAP ratio, and **(H)** gm130 expression (n = 8). Bars represent mean ± SEM. All data was analyzed using two-way ANOVAs with Tukey post hoc analyses to evaluate NEI and YAP effects. For streams, *’s denote the significance level for YAP effects. Time plots of migration characteristics (n ≥ 6): **(I)** net velocity, **(J)** order parameter, **(K)** percentage of backtracking vectors, and **(L)** speed for α-cat KD monolayers on 2 kPa polyacrylamide gel. Lines represent mean ± SEM. Scale bars: 50 µm.

By itself, α-catenin knockdown (α-cat KD) destabilized intercellular adhesions and yielded a mesenchymal phenotype (Fig. 8A, D). Cells at the leading edge best epitomized mesenchymal characteristics, exhibiting more morphological polarization, aided by their separation from the cells behind. Cells residing closer to the monolayer center remained bunched due to contact inhibition. However, cell-cell adhesion there was still low. NEI generally caused re-epithelialization of α-cat KD cells, during which cells lost front-back polarity, formed discrete colonies, and dissolved leader-follower relationships (Fig. 8A, D). However, while NEI caused a nearly 3 times increase in intercellular adhesions (Fig. 8C) and promoted nuclear accumulation of IκBα (Fig. 8B), p120 expression was markedly lower than WT for all conditions. These results suggest that interrupting force propagation through intercellular adhesions decreases the epithelial features of NEI’s concurrent E-M phenotype.

Without this cooperation between intercellular adhesions and intracellular forces intrinsic to migrating epithelia, cells treated with vehicle migrated as a mesenchymal collective (Theveneau and Mayor, 2013). α-cat KD cells migrated with lower speed and order, exhibiting more backtracking and thus lower net velocity relative to WT (Fig 8I-J). However, likely arising from contact inhibition, these cells developed order and net velocity over time (Fig. 8I, J). Meanwhile, in removing the cooperative force balance, data suggest we also severed the link required to sustain NEI’s concurrent E-M state. Without force propagation between cells, the effect of NEI on migration was time-dependent across stiffnesses. Initially, NEI elevated net velocity. From 0-4 h on 0.5 and 50 kPa, net velocities were higher than vehicle control. On 2 kPa, this initial phase extended to 8 h. After that phase, velocities decreased and approached zero by 24 h (Fig. S6 A-C). Differences in order underlay the differences in velocity. During the initial time points, cells migrated with slightly higher order and speed (Fig. 8L) and thus achieved higher collective velocity. However, while higher speeds endured for the time-lapse duration, there was progressive disarray and loss of net velocity. Cell backtracking mirrored these changes (Fig. 8K). Overall, findings suggest that α-catenin knockdown exacerbates the loss of order during NEI to facilitate the arrest of net migration. In other words, intercellular adhesion reinforcement and YAP may jointly enhance cohesion to slow the loss of order during NEI. Outcomes from removing α-catenin further imply an unusual partnership paramount to the concurrent E-M phenotype, where better intercellular adhesion facilitates the propagation of higher forces through the collective to sustain the simultaneous E-M state.

## Discussion

Classically, epithelial (E) and mesenchymal (M) phenotypes exist along a continuum, where cells traverse from E to M via the gradual loss of epithelial characteristics and the accompanying gain of mesenchymal ones. Given the diversity of EMT programs and the array of TFs and RNA regulators associated with protein regulation, groups have encouraged describing E-M phenotypes as functional cellular changes (Yang et al., 2020). Moreover, greater appreciation for E-M phenotypes as an aggregate outcome from the activity of all EMT-related proteins has emphasized that single biological markers are not sufficient to categorize cellular states. Together, these statements pose a compelling question about the interaction between EMT-promoting and -inhibiting transcription factors during the development of cell phenotypes: do these proteins inherently compete, cooperate, or cancel? Here, we use Leptomycin B, a CRM1-based inhibitor of nuclear export, to explore this concept. We demonstrate that, across substrate stiffnesses, MCF10A collectives simultaneously develop mechanoactive features and reinforce intercellular adhesions during NEI. Within the collective, both leaders and followers exhibit p120-mediated adhesion strengthening. Meanwhile, YAP, pMLC, and gm130 expression globally increase. Leader cells exaggerate protrusions, and those protrusions demonstrate higher coherency of F-actin. Ultimately, within the collective, we show that cells manifest atypical E-M states, in which elements at opposite ends of the conventional spectrum coexist (Fig. 9A). Interestingly, these results suggest the classical view that epithelial and mesenchymal phenotypes are antagonistic – and accordingly exist along a continuum – may alone be insufficient. Instead, our findings indicate that opposing transcriptional programs can simultaneously give rise to highly epithelial and highly mesenchymal characteristics, which we refer to as concurrent E-M.

**Figure 9.**
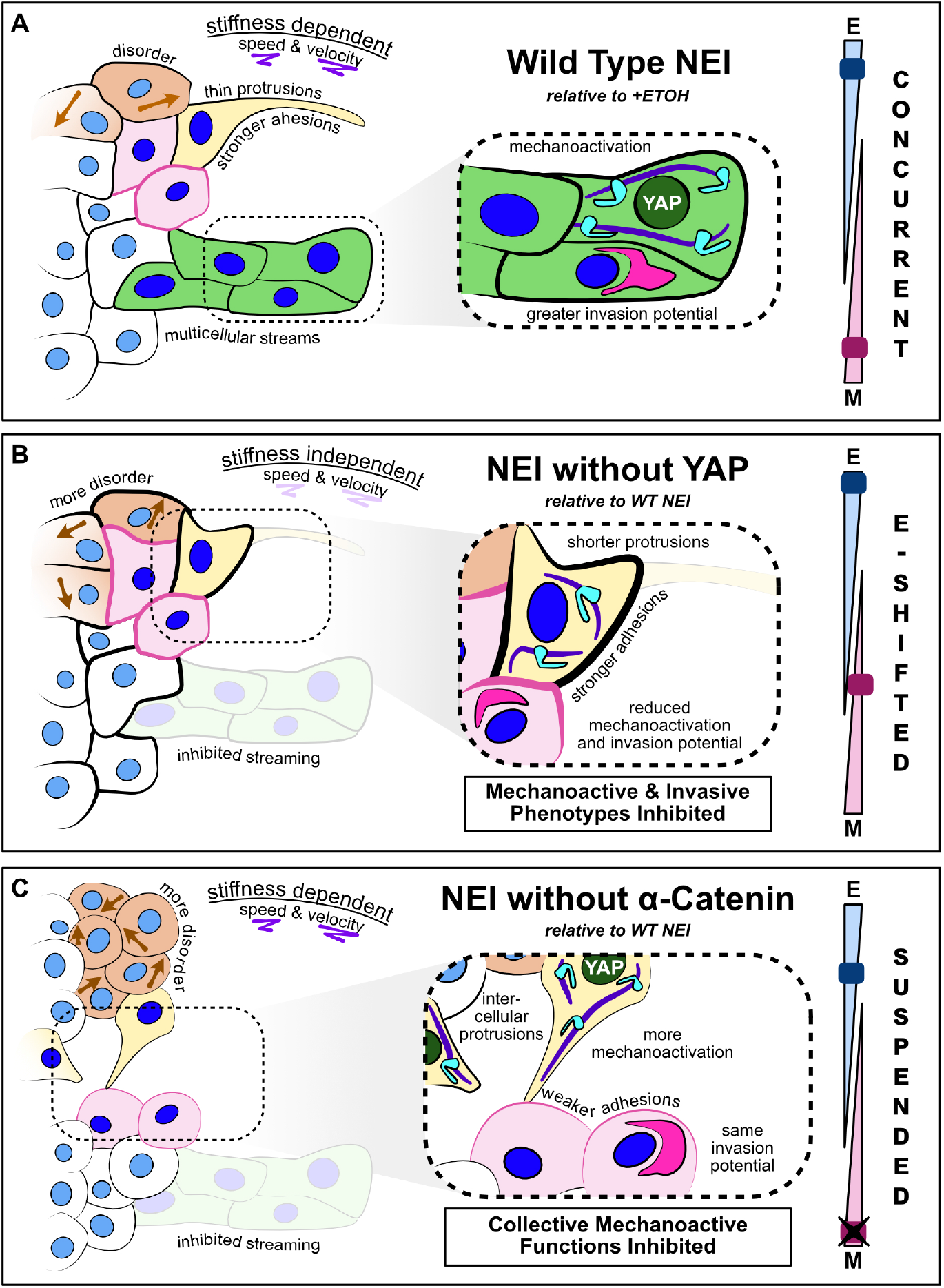
Summary schematic depicting NEI outcomes for WT, shYAP, and α-catenin knockdown cells. **(A)** Relative to vehicle, WT MCF10A exhibit stronger intercellular adhesions, together with higher mechanoactivation and invasion potential. Cells exhibit thin protrusive morphology as single cells and form more multicellular streams at the leading edge. We propose this is a concurrent epithelial-mesenchymal (E-M) state. NEI disrupts collective migration, causing stiffness-dependent changes in speed and velocity, and overall migratory disorder. **(B)** Relative to WT NEI, NEI for shYAP yields stronger intercellular adhesions while lowering mechanoactivation and invasion potential. Cells form shorter protrusions and are unable to form multicellular streams. We propose this is an E-shifted concurrent state. **(C)** Relative to WT NEI, NEI for α-cat KD MCF10A yields weaker intercellular adhesions and more mechanoactivation. Cells protrude to form more intercellular connections, but collective mechanoactive function is inhibited. Given the strong mechanoactive signatures, but low mechanoactive behavior, we propose that α-cat KD suspends the concurrent E-M state.

We further demonstrate that within a simultaneous state, epithelial and mesenchymal characteristics individually vary. We show that excluding YAP from the NEI-affected cargos attenuates some mesenchymal phenotypic components while strengthening epithelial ones (Fig. 9B). This result suggests a conceptual model where the strength of epithelial and mesenchymal features can together traverse a range of distances from the classical center-point. Our experiments additionally revealed that knockdown of α-catenin suspended the concurrent phenotype (Fig. 9C). Thus, the manifestation of simultaneous E-M characteristics may hinge on cooperation between intercellular adhesion and intracellular forces. Given α-catenin’s role in recruiting F-actin to adherens junctions and instituting tension across cell-cell contacts, this is perhaps not surprising. Interestingly, it suggests that this phenomenon could resolve a problem fundamental to collective migration: how follower cells maintain high adhesion to neighboring cells while simultaneously extending cryptic protrusions to assist grouped migration (Lu and Lu, 2020). In contrast to the disordered migratory patterns of the concurrent phenotype observed after NEI, under normal conditions, followers sustain high cooperation between epithelial and mesenchymal features. Here, epithelial and mesenchymal characteristics extended far from the phenotypic center point. Meanwhile, typical follower cells manifest moderately high adhesion and moderately high protrusive ability (Desai et al., 2013; Qin et al., 2021). Therefore, it is possible that the degree of cooperation during concurrent E-M may ultimately depend on the distance of either E-M component from the phenotypic center.

Beyond disrupting this force balance, NEI changed conventional leader-follower relationships. Typically, leader cells exhibit lower intercellular adhesion, and previous studies suggest this is required to conduct their role as the primary drivers of migration (Qin et al., 2021; Matsuzawa et al., 2018), where they polarize front-back and generate most of the traction to propel the collective (Reffay et al., 2014; Trepat and Sahai, 2018). Here, we show that NEI maintained the gradient of intercellular adhesion, such that adhesions between leaders remained weaker than those between followers. However, the increase in cell-cell adhesion by itself is presumably sufficient to disrupt the function of leader cells. In general, higher cell-cell adhesion predisposes cells to repolarization cues from neighbors and reduces the contribution of polarization stemming from mechanosensitive adhesion-based promotion of locomotion (Desai et al., 2013; Qin et al., 2021). Without appropriate polarization signals, leaders would not be able to effectively coordinate movement of the group. Therefore, changes in the intercellular adhesion strength for leaders could itself contribute to the observed disordered migration. While NEI changed the intercellular adhesion of leaders and followers, it also disrupted their characteristic patterns of mechanoactivation. In typical epithelia, leader cells exhibit higher mechanoactivation and a more mesenchymal phenotype. Enrichment of these features is greatest in cells at the leading edge that actively drive the formation of finger-like streams (Sarker et al., 2019). In our experiments, these mesenchymally-enriched leaders appeared at a higher frequency and with exaggerated morphology after NEI, as if competing to guide migration of the group. This contest between leaders could also have contributed to the disorder. Meanwhile, competition with followers may have caused further disarray. After NEI, follower cells displayed similar YAP and pMLC levels to leaders, which suggests a greater capacity for follower-generated traction. Indeed, in control monolayers we observed tractions localized primarily to the leading edge (Fig. S7A). Meanwhile, NEI yielded tractions extending farther into the monolayer core; and overall traction magnitudes were higher compared to control (Fig. S7B). Altogether, these changes in mechanoactivation imply a disruption to group dynamics, where more cells are individually driving migration. From a broader standpoint, these results emphasize that epithelial and mesenchymal characteristics can exist in high degrees concurrently but not necessarily cooperatively.

### Ideas and Speculation

> At a basic level, the cohesion of epithelial collectives derives from a balance between intercellular adhesion strength and cell-generated traction forces (Trepat and Sahai, 2018). After NEI, the migratory order of epithelia was generally low; however, it appeared to exhibit a biphasic relationship with substrate stiffness (Fig. S8). We observed that cell-cell adhesion strength was lowest on 2 kPa and higher on both the softer and stiffer substrates (Fig. S8A). Meanwhile, mechanoactivation was high across stiffnesses (Fig. S8B-E). These distinct outcomes suggest that NEI may disrupt the balance between intercellular adhesion and cell traction required for collective cohesion. As an illustration of this idea, higher percentage losses in migration speed, velocity, order, and polarization (Fig. S8F-K) correlated with lower adhesion strength (i.e., 2 kPa). This biphasic trend across migration parameters extended to stream formation. The higher disorder on 2 kPa implied that epithelia could encounter greater leading-edge instability, and 2 kPa indeed coincided with a higher protrusion rate of multicellular streams (Fig. S8J). However, despite this increase in stream formation, lower cell cohesion limited the number of cells that successfully coordinated their velocities to establish streams (Fig. S8K). Altogether, the balance between intercellular adhesion and mechanoactivation was most uneven on 2 kPa, and the disorder was worst there. Therefore, these findings suggest that, while NEI enhances both epithelial and mesenchymal traits, cells may not develop strong enough intercellular adhesions to balance cellular tractions, and this imbalance may ultimately contribute to the observed migratory disorder.

The presented findings fill an important gap in the biophysical understanding of nuclear export inhibition, connecting the subcellular trapping of competing E-M factors to disruptive phenotypic changes in healthy epithelia. This updated biophysical understanding could inform additional studies regarding the clinical translation of NEI. Clinically, NEI has promising potential to treat cancer – largely for its ability to re-localize escaped tumor suppressors to the nucleus (Zhong et al., 2014; Kashyap et al., 2016a; Kashyap et al., 2016b; Galinski et al., 2021; Gravina et al., 2014; Gravina et al., 2017; Turner et al., 2014). However, because EMT is a driver for cancer metastasis and changes the function of healthy cells, our findings of mechanoactivation could affect the efficacy and specificity of these treatments. Interestingly, multiple studies investigating NEI in this clinical context have suggested that NEI causes EMT reversal in cancer cells, with inhibition of NFKB by IκBα playing a significant role (Kashyap et al., 2016a; Kashyap et al., 2016b; Galinski et al., 2021). However, such studies have not yet surveyed mechanoactive and mesenchymal markers, nor have they investigated cellular outcomes on substrates better approximating the stiffnesses of physiological tissues. Given the stiffness-induced alteration of nuclear pore architecture and the associated change in transport selectivity based on cargo molecular weights and stabilities (Elosegui-Artola et al., 2017), the high stiffness of standard tissue culture plastic could significantly alter the emergence of cell phenotypes. It may also mask some of the outcomes observed here. Ultimately, whether the outcomes we observe here manifest similarly in clinical versions of NEI, and more pointedly, whether mechanoactivation could antagonize therapeutic effects remains to be seen.

To put these results in an appropriate context, we acknowledge several limitations of our study. Our experiments sought to determine how EMT-promoting and –inhibiting proteins interact during the development of cell phenotypes. We used NEI to implement the complex situation where competing EMT-related proteins could interact. However, CRM1 has ∼241 cargos and therefore affects many proteins beyond the explicit EMT scope (Turner et al., 2014; Xu et al., 2012). To account for this nonspecificity, we frame our conclusions as phenotypic outcomes of NEI. Still, we believe the outcomes themselves retain significance without the identification of protein-specific mechanisms. To build on this work, future studies may seek to incorporate newer techniques that enable better control over the localization of individual proteins. It is also important to point out that the outcomes we observe from NEI are not generalizable, as there are a variety of methods to induce NEI. Here, we used leptomycin B, which causes permanent inhibition of CRM1-based nuclear export. Similar drugs, e.g., Selective Inhibitors of Nuclear Export (SINEs), only transiently inhibit the CRM1 nuclear export receptor (Senapedis et al., 2014). Their reversible binding to CRM1 would likely engender rate-dependent differences in migration characteristics and cell phenotypes. This constitutes an interesting and clinically-significant question for future investigations. Another point of limitation arises from the method we use to shift the concurrent E-M phenotype. Here, we silence YAP to determine how its exclusion from NEI affects epithelial and mesenchymal characteristics. As a byproduct of this silencing, we remove the cytoplasmic role of YAP, which overlaps with NFκB regulation. Thus, conclusions pertaining to YAP silencing must not be considered a sole outcome of inhibiting YAP activation but its absence altogether (Wang et al., 2020). Indeed, the results are consistent with both YAP de-activation (increased intercellular adhesion and reduced mechanoactive characteristics) and YAP silencing (NFκB activation through IκBα degradation). Throughout the discussion of data, we refer to these results transparently as outcomes of YAP silencing. Still, we reason that YAP silencing is ultimately instructive in this context, as it reveals an adjustable range for concurrent E-M characteristics. These limitations, taken together with the presented results, pave additional avenues for investigation.

In sum, here we show that NEI promotes simultaneous intercellular adhesion strengthening and mechanoactivation, which we propose constitutes a concurrent E-M state in which highly epithelial and highly mesenchymal characteristics coexist. This concurrent manifestation complicates the classical interpretation of phenotypes and, ultimately, further encourages a more comprehensive assessment of E-M cellular features as a basis for phenotypic categorization. We show that the simultaneous phenotype here hinges on cooperation between intercellular adhesion and intracellular force and that it has an adjustable range. Given this flexibility, we hypothesize that some simultaneous E-M states may manifest cooperatively, such as in follower cells during collective migration. Moreover, as a point for future investigation, we speculate that these biophysical outcomes may translate clinically, where the observed mechanoactive effects and stiffness-dependent differences could influence the efficacy and specificity of NEI-centered treatments.

## Acknowledgements

This work was supported by the NIH/NIGMS MIRA (R35GM128764) grant to AP.

## Author Contributions

CMK, CL, and AP conceived the project and designed experiments. CMK and CL performed experiments and analyzed data. CMK interpreted findings, made figures, and wrote the manuscript. AP edited the manuscript, supervised the project, and acquired funding.

## Materials and Methods

### Key resources table

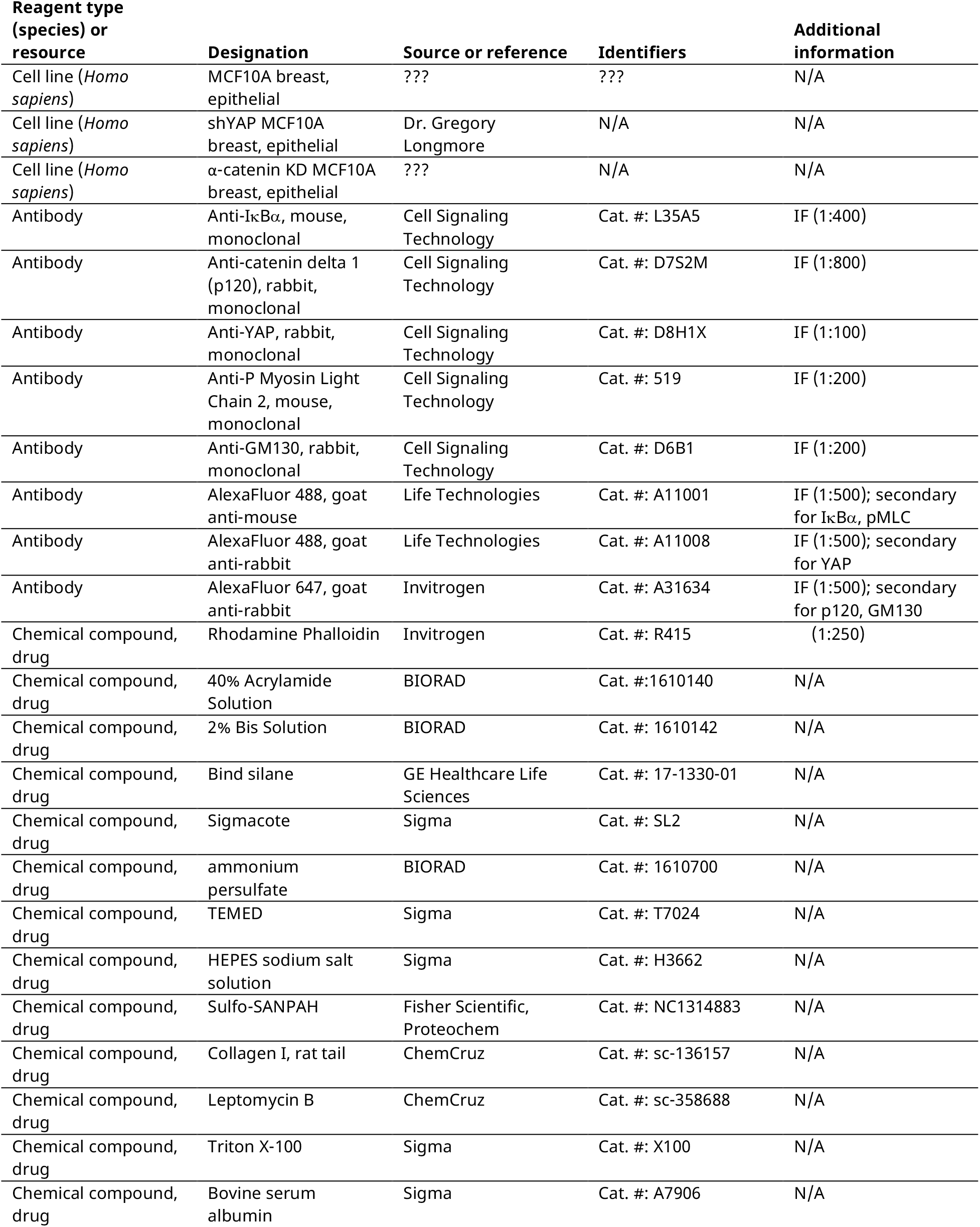

### Polyacrylamide gel synthesis and collagen coating

Polyacrylamide gels were synthesized via free-radical polymerization, according to established protocols (Fischer et al., 2012). Precursor solutions were formulated from acrylamide, bis-acrylamide, and ultrapure water. To yield gel stiffnesses of 0.5, 2, and 50 kPa, respective acrylamide and bis-acrylamide percentages of 4%/0.2%, 5%/.228%, 12%/.6% were used. Gels were attached to glass coverslips for immunofluorescent analyses and to glass-bottomed plates for time-lapse imaging. To facilitate gel attachment, these glass surfaces were activated by treatment with plasma and bind silane solution (94.7% ethanol, 5% acetic acid, and 0.3% bind silane for 10 min). After treatment, surfaces were rinsed with ethanol and air-dried. Precursor polyacrylamide solutions were polymerized by adding 10% ammonium sulfate and N,N,N’,N’-tetramethylethylene (TEMED) at respective ratios of 1:200 and 1:2000 v/v. For immunofluorescent experiments, these final solutions were immediately dispensed onto Sigmacote-treated microscope slides and covered with plasma-activated glass coverslips; meanwhile, for time-lapse experiments, final solutions were dispensed onto plasma-activated glass-bottomed plates and covered with Sigmacote-treated glass coverslips. Solutions were allowed to polymerize for 30 min, then Sigmacote-treated surfaces were removed to expose one face of the formed gels. Gels were functionalized by 10 min incubation with Sulfo-SANPAH solution (50 mM HEPES buffer and 0.5 mg/mL Sulfo-SANPAH in dH_2_ O) under 365 nm UV. After rinsing, collagen type I solution (0.5 mg/mL in PBS) was applied, and gels were stored at 4 °C overnight.

### Cell culture and colony seeding

MCF10A cell lines were cultured at 37°C and 5% CO_2_ in DMEM/F12 supplemented with 5% horse serum, 20 ng/ml epidermal growth factor, 0.5 mg/ml hydrocortisone, 100 ng/ml cholera toxin, 10 µg/ml insulin, and 0.2% Normocin. Media was changed every 3 days during cell expansion. To prepare for cell colony seeding, collagen-coated polyacrylamide gels were rinsed gently with PBS and allowed to dry for 20 min. A volume of 5 µL, containing 18,000 cells, was then seeded at the center of each gel. Seeded cells were incubated for 24 h to facilitate attachment and acclimatization to gel stiffness. After this incubation period, cells were treated with 100 nM leptomycin B (LMB, constituted in 100% ethanol) or vehicle. Treated cell colonies were then used for immunofluorescent staining or time-lapse imaging.

### Immunofluorescence and confocal microscopy

Treated cells were cultured for 24 h, then were fixed with 4 % paraformaldehyde for 15 min. Fixed cells were rinsed and stored at 4 °C until immunostaining. For cell membrane permeabilization, wells were incubated with 0.3 % Triton X-100 in 2 % w/v BSA at room temperature for 10 min. Then wells were blocked using 2 % BSA in PBS, with overnight incubation at 4 °C. Primary antibodies were incubated overnight at 4 °C. After rinsing, secondary antibodies were also incubated overnight at 4 °C. After secondary incubation, wells were rinsed and incubated with DAPI and phalloidin (if applicable) for 45 min. After final rinses, gels were coverslipped with mounting medium, and allowed to dry overnight. Immunostained cells were imaged at 40x on a confocal microscope. To facilitate sample comparability, optimal capture settings were determined for each antigen based on the highest observed signals, then those settings were maintained for the duration of imaging.

### Time-lapse microscopy

Time-lapse imaging was performed using a Zeiss AxioObserver Z1 microscope. Phase contrast images were acquired at 10 min intervals for a 24 h period using a 10x objective. For the duration of imaging, cell colonies were maintained at 37°C and 5% CO_2_ using an attached incubation chamber.

### Quantitative image analysis

Unless otherwise stated, results for each condition (vehicle, LMB) reflect 8 FOVs from 2 biological replicates. Collected phase contrast time-lapse images were analyzed for the presence of multicellular streams. Inclusion criteria for streams were: (1) protrusion of 2 or more cells in succession from the monolayer baseline and (2) a protrusion length greater than twice its width (Sarker et al., 2019). To account for differences in image orientation, the number of streams counted over 24 h was normalized to the baseline length of the migrating cell colony. All protein expression was measured in ImageJ. Nucleocytoplasmic (N:C) ratio was calculated as the nuclear intensity integrated over the nuclear area divided by the cytoplasmic intensity integrated over the cytoplasmic area. For YAP, N:C ratio was measured in at least 55 leader cells and 220 follower cells per condition. p120 expression (n > 39 leader cells and 55 follower cells from at least 6 FOVs reflecting 2 biological replicates per condition) was quantified as the average signal intensity integrated over a 10 um line, where the line lay perpendicular to the junction and its midpoint rested at the junction center. 3 measurements per cell-cell junction were averaged to define a single junction intensity. For α-cat KD, adhesions in cells treated with vehicle were mostly undetectable, so p120 expression was considered an unreliable measure. Instead, differences between groups were assessed via the number of visible adhesions. pMLC and gm130 expression were measured as intensity within the monolayer, and were normalized to DAPI signal. Actin coherency was measured using the OrientationJ plugin, where n reflected the number of coherency measurements. Protrusions from the monolayer baseline were outlined manually, and the vector field was extracting using a grid size of 10 and an α value of 2. Because α-cat KD cells segregated into distinct colonies without clear polarization, coherency was measured for all cells. To assess Golgi orientation, the angle between the direction of migration and the line connecting a particular cell’s nuclear centroid to its Golgi centroid was measured (n > 50 leader cells and 250 follower cells from at least 8 FOVs reflecting 2 biological replicates per condition). A polarized Golgi was considered to be one where this angle was less than 60°, as defined previously (Mason et al., 2019).

### Particle image velocimetry

Particle image velocimetry (PIV) was used to extract spatiotemporal velocity profiles from time-lapse images of migrating cell colonies. This analysis was performed using the MATLAB PIVlab package. The velocity field was calculated using 3 passes, with decreasing window sizes of 64, 32, and 16 pixels. All migration characteristics and heat maps were computed using MATLAB code (n > 6 FOV from 2 biological replicates per condition). The degree of cell cohesion was measured using order parameter, defined as the cosine of the angle between a single velocity vector and the average velocity vector for the respective time point. Meanwhile, when the value of that angle was greater or less than 90°, the vector was counted as backtracking.

### Traction Force Microscopy

To measure gel deformations, 1 µm fluorescent polystyrene beads were mixed with 0.5 kPa polyacrylamide solution at a concentration of 20 million beads per mL gel solution. Images of beads at the gel surface (<10 µm depth) were captured during migration with vehicle or LMB. An image of the relaxed beads, taken after cells were detached from gels using 0.5% trypsin, was used for reference to determine bead displacements. Bead displacements and cell-generated tractions were calculated using the Particle Image Velocimetry and Traction Force Microscopy ImageJ plugins, as described previously (Tseng et al., 2012). Heat maps were generated using MATLAB code.

### Statistical analysis

Bar and line plots are presented as the mean ± SEM. Statistical significance was determined using two-way ANOVA with Tukey post hoc comparisons. For tractions, significance was assessed using two-tailed t-test. Differences were considered significant for *p* < .05.

