## Supplemental Information for "Nuclear export inhibition jumbles epithelial-mesenchymal states and gives rise to migratory disorder in healthy epithelia"

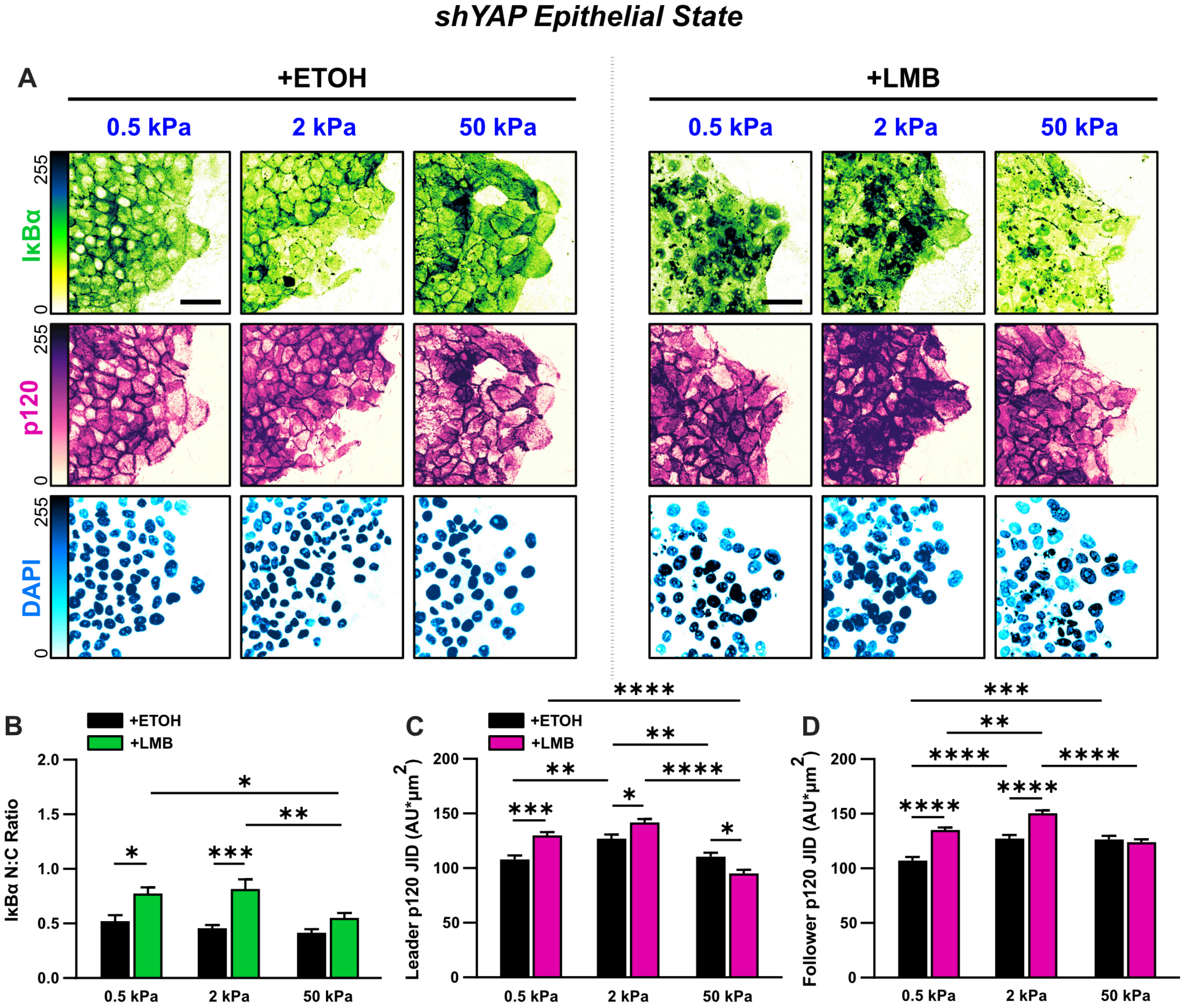


**Figure S1. Epithelial characteristics for shYAP cells for all stiffnesses.**

**(A)** Representative images for epithelial characteristics of shYAP MCF10A on 0.5, 2, and 50 kPa polyacrylamide gels. Monolayers were treated with ethanol (ETOH) as vehicle or leptomycin B (LMB) for NEI. Images depict nucleocytoplasmic localization of IκBα (top), p120 expression (middle), and DAPI nuclear signal (bottom). **(B)** shYAP nucleocytoplasmic (N:C) ratio for IκBα (n > 6). **(C)** Leader and **(D)** **f**ollower changes in p120 junction integrated density (JID) for shYAP cells (n = 55 leaders and 55 followers). Data was analyzed using two-way ANOVA with Tukey post hoc analyses to evaluate NEI and stiffness differences. Bars represent mean ± SEM. Scale bars: 50 μm.

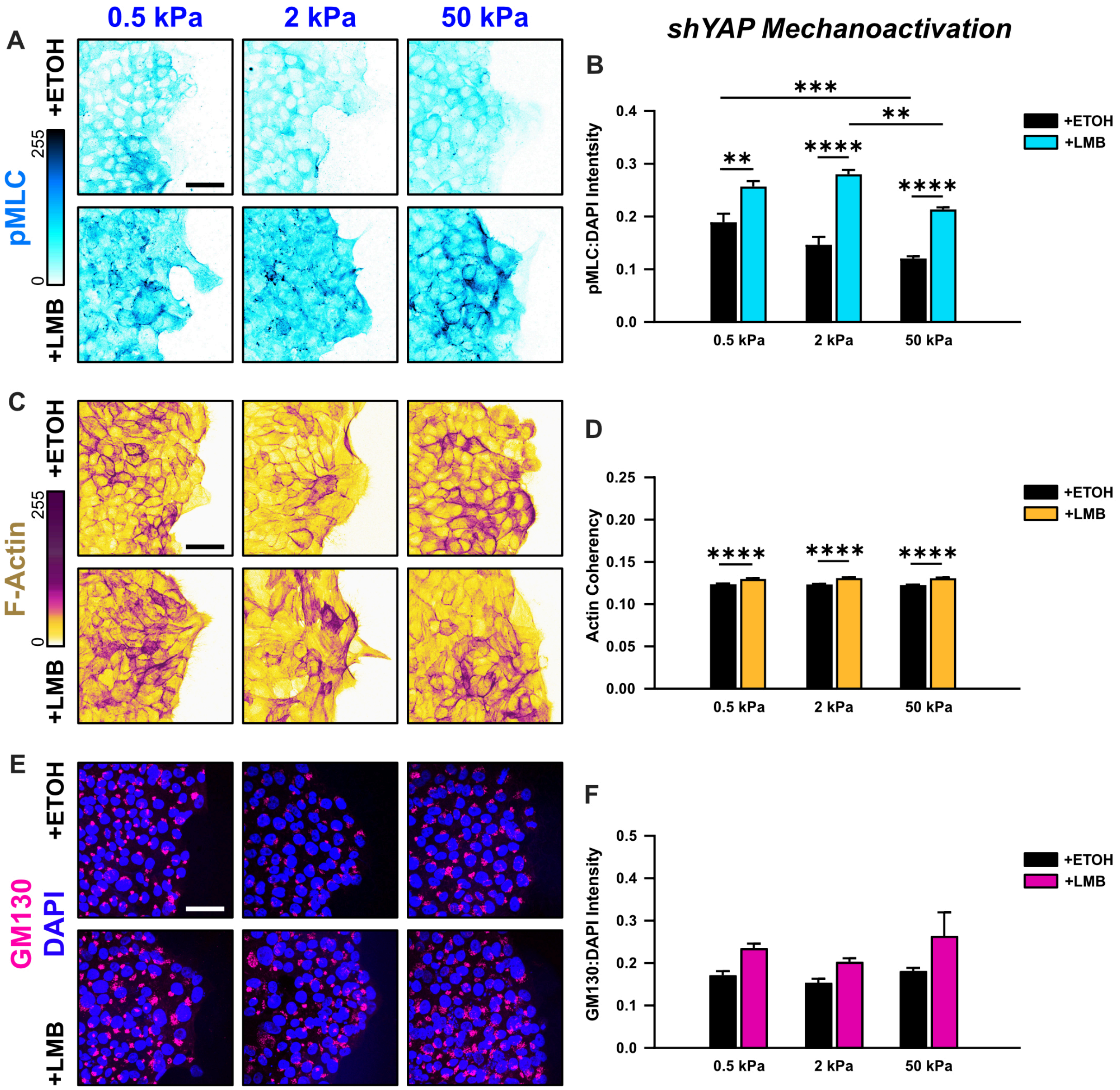


**Figure S2. Mechanoactive characteristics for shYAP cells for all stiffnesses.**

**(A)** Representative images of shYAP cells for pMLC expression across stiffnesses. **(B)** Quantification of pMLC intensity (n > 6). **(C)** Representative images of shYAP cells for F-actin across stiffnesses. **(D)** Quantification of actin coherency (n > 1000). **(E)** Representative images of shYAP cells for gm130 across stiffnesses. **(F)** Quantification of gm130 intensity (n = 8). Data was analyzed using two-way ANOVAs to evaluate with Tukey post hoc analyses effects from NEI and stiffness. Bars represent mean ± SEM. Scale bars: 50 μm.

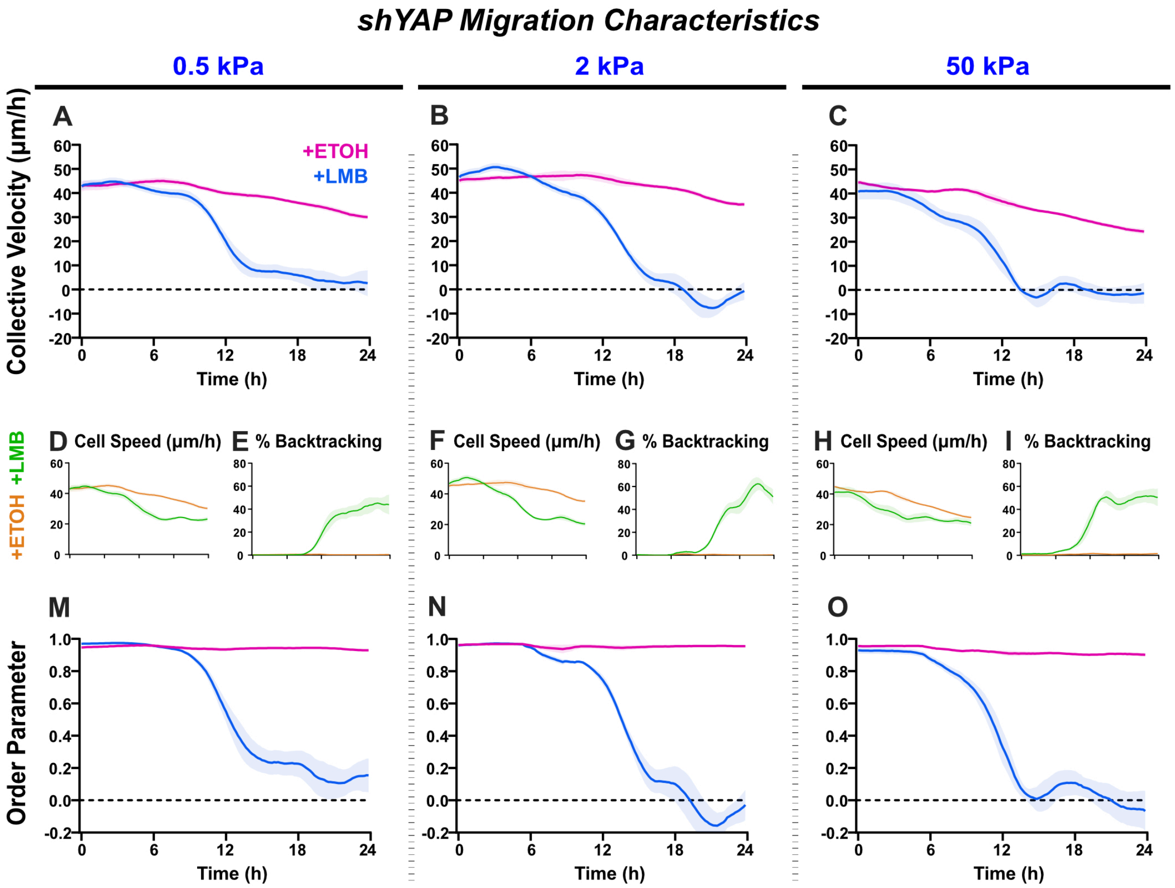


**Figure S3. shYAP migration characteristics across stiffnesses.**

Net migration velocity for shYAP on **(A)** 0.5 kPa, **(B)** 2 kPa, and **(C)** 50 kPa. Accompanying speeds **(D, F, H)**, percentages of backtracking vectors **(E, G, I),** and order parameters **(M, N, O)**.

### *α-cat KD Epithelial State*

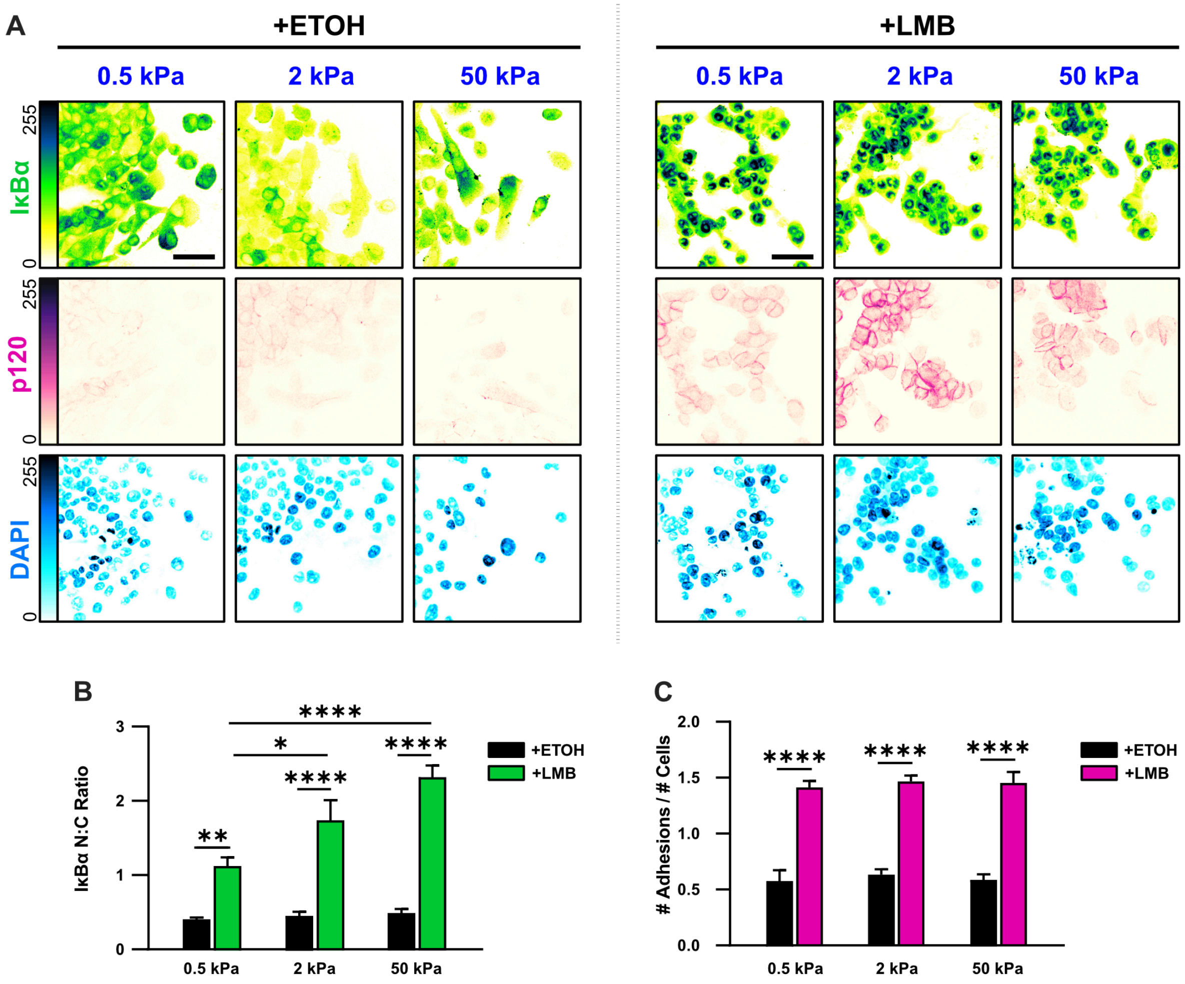


**Figure S4. Epithelial characteristics for α-cat KD cells for all stiffnesses.**

**(A)** Representative images for epithelial characteristics of α-cat KD MCF10A on 0.5, 2, and 50 kPa polyacrylamide gels. Monolayers were treated with ethanol (ETOH) as vehicle or leptomycin B (LMB) for NEI. Images depict nucleocytoplasmic localization of IκBα (top), p120 expression (middle), and DAPI nuclear signal (bottom). **(B)** α-cat KD nucleocytoplasmic (N:C) ratios for IκBα (n > 6). **(C)** Differences in the number of discernible p120-marked junctions (n > 6). Data was analyzed using a two-way ANOVA with Tukey post hoc analyses to evaluate NEI and stiffness effects. Bars represent mean ± SEM. Scale bar: 50 μm.

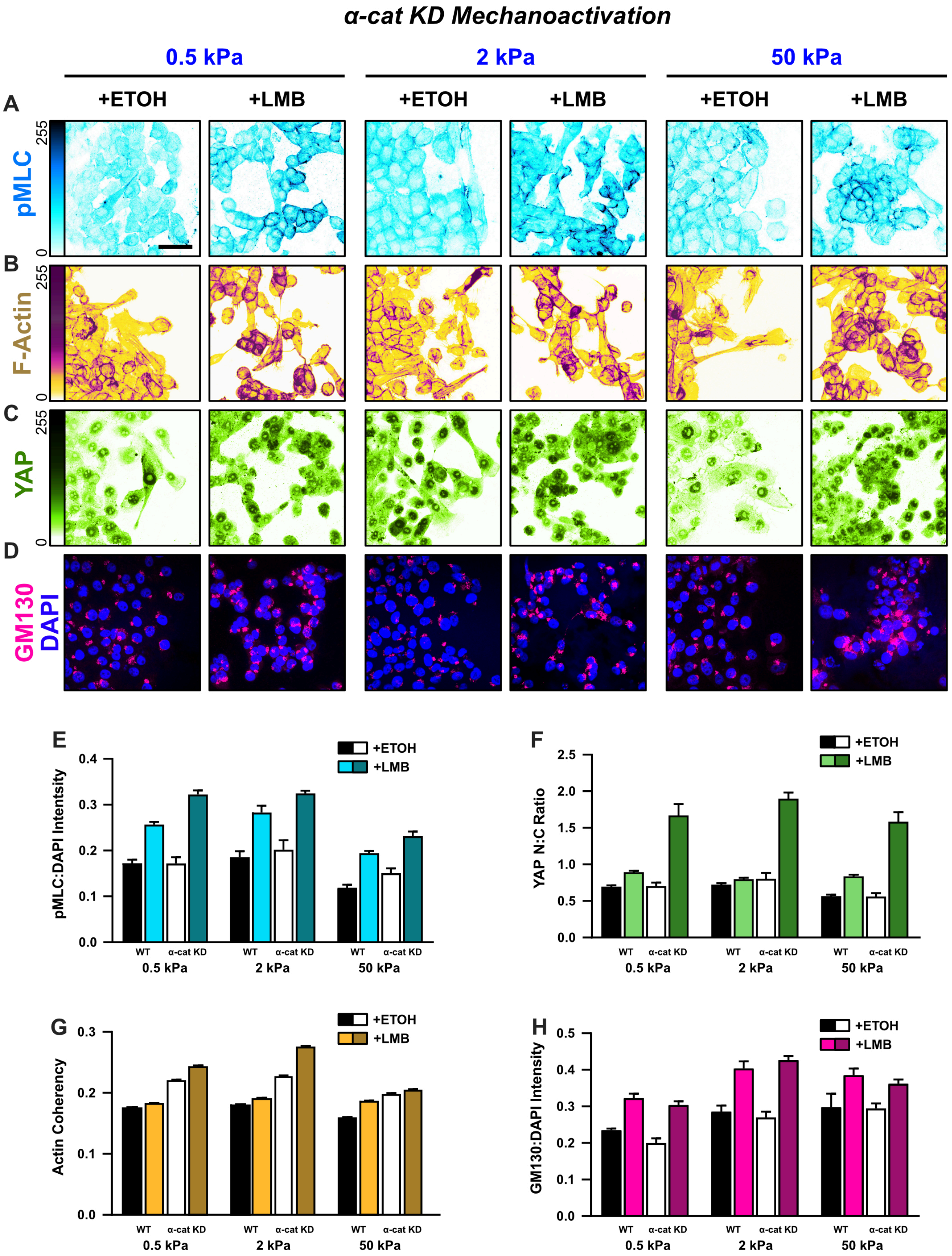


**Figure S5. Mechanoactive characteristics for α-cat KD cells for all stiffnesses.**

Representative images of α-cat KD cells for **(A)** pMLC, **(B)** F-actin, **(C)** YAP, and **(D)** gm130 across stiffnesses. **(E)** α-cat KD quantification of pMLC expression (n = 8), **(F)** YAP nucleocytoplasmic (N:C) ratio, **(G)** actin coherency (n > 1000), and **(H)** gm130 expression (n = 8) across stiffnesses. Data was analyzed using two-way ANOVAs with Tukey post hoc analyses to evaluate effects from NEI and stiffness. Bars represent mean ± SEM. Scale bar: 50 μm.

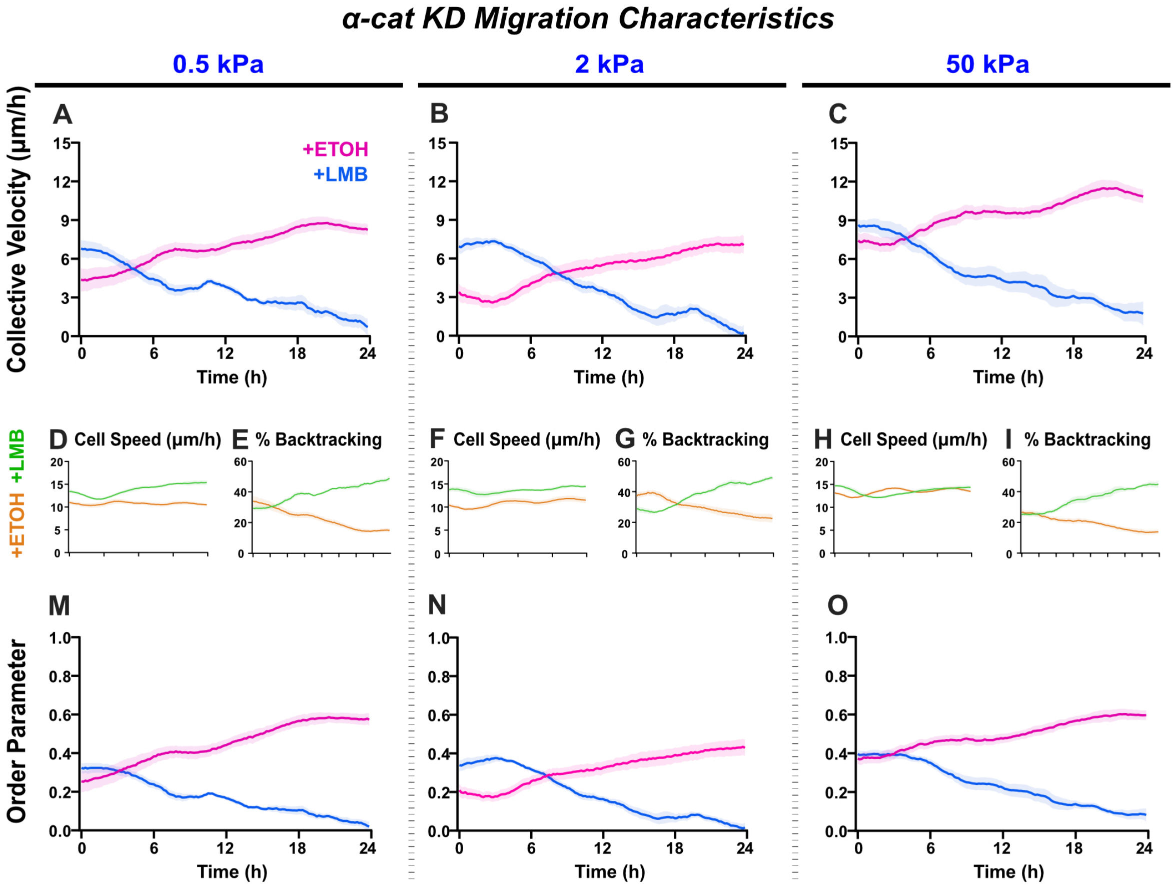


**Figure S6. α-cat KD migration characteristics across stiffnesses.**

Net migration velocity for shYAP on **(A)** 0.5 kPa, **(B)** 2 kPa, and **(C)** 50 kPa. Accompanying speeds **(D, F, H)**, percentages of backtracking vectors **(E, G, I),** and order parameters **(M, N, O)**.

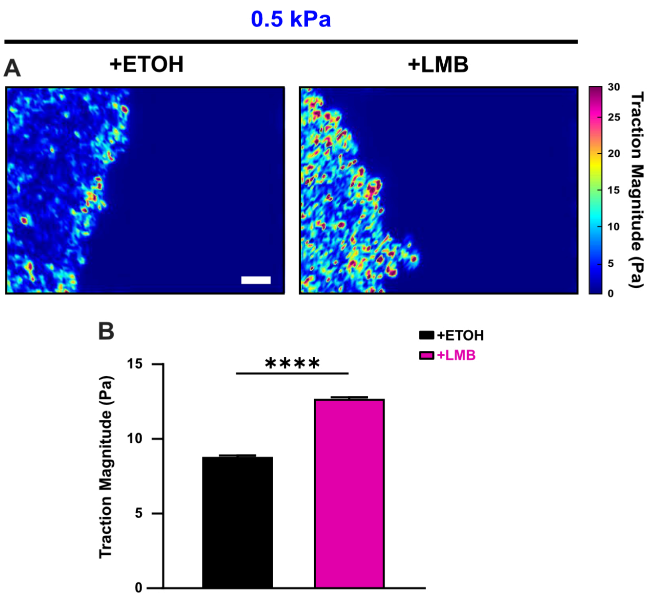


**Figure S7. NEI increases cell-generated tractions.**
**(A)** Traction heat maps and **(B)** quantified tractions for vehicle (n = 11) and NEI (n = 15) –treated cells on 0.5 kPa polyacrylamide gel. Differences were assessed via two-tailed t-test. Bars represent mean ± SEM. Scale bar: 100 μm.

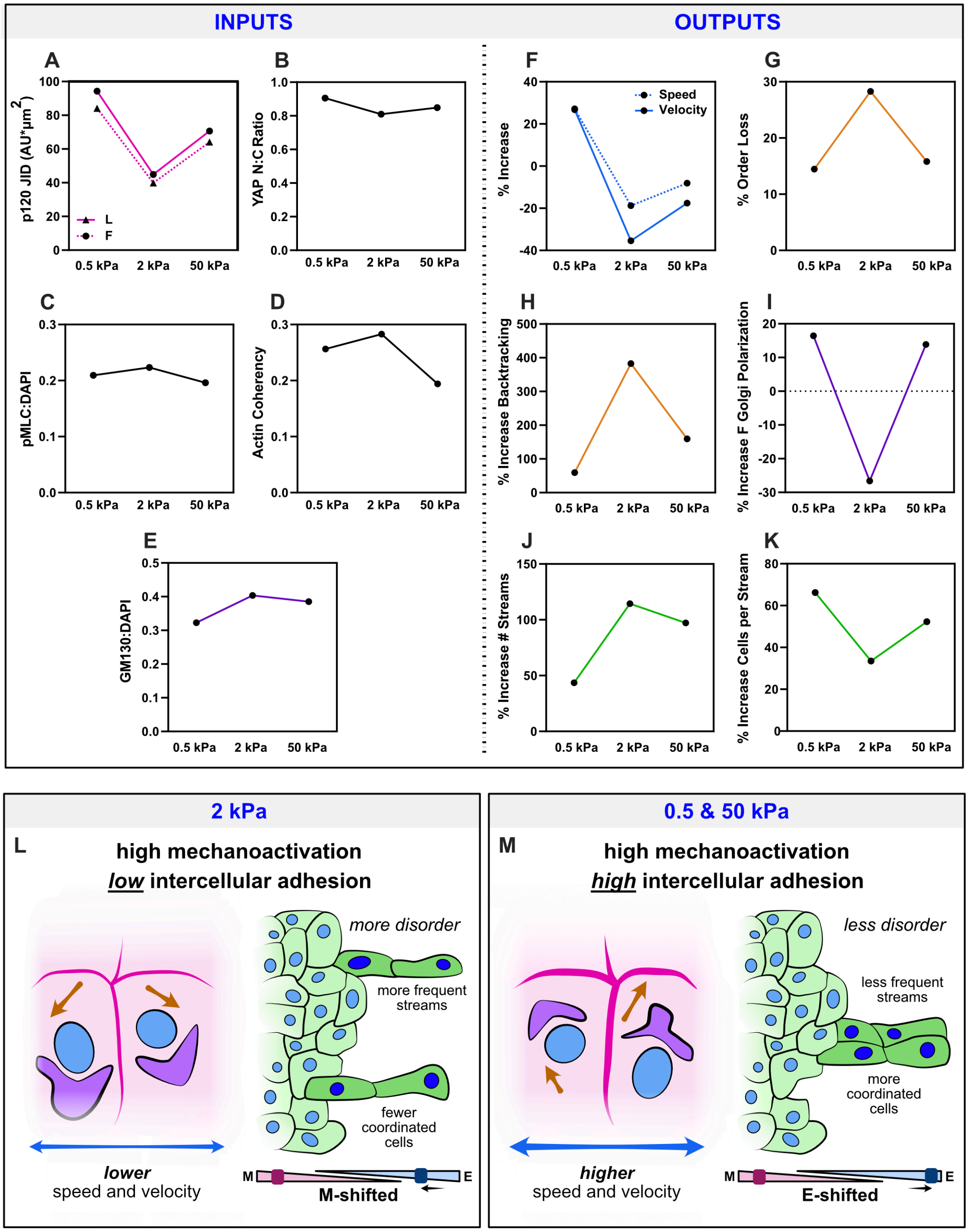


**Figure S8. Outcomes from WT NEI are biphasic with substrate stiffness and may derive from shifts in the balance between intercellular adhesion and mechanoactivation.**

Average measures of **(A)** p120 adhesion strength, **(B)** YAP nucleocytoplasmic (N:C) ratio, **(C)** pMLC expression, **(D)** actin coherency, and **(E)** gm130 expression for each stiffness. Accompanying percentage changes for **(F)** speed and velocity, **(G)** order parameter, **(H)** backtracking, **(I)** follower Golgi polarization, **(J)** stream formation, and **(K)** the number of cells per stream. **(L)** Schematic illustrating relationships on 2 kPa, where lower adhesion strength correlates with the overall higher disorder. **(M)** Schematic illustrating relationships on 0.5 and 50 kPa, where higher adhesion strength correlates with the overall lower disorder.
